# FERRET: Framework to Evaluate Robustness in Regulatory Networks Using Heterogeneous Cell Types

**DOI:** 10.64898/2026.07.21.739795

**Authors:** Tara Eicher, John Quackenbush

**Affiliations:** Department of Biostatistics, Harvard T.H. Chan School of Public Health, Boston, MA, 02115, USA; Channing Division of Network Medicine, Brigham and Women’s Hospital, Boston, MA, 02115, USA

## Abstract

Techniques for evaluating gene regulatory network (GRN) inference methods typically focus on recovering small ground-truth networks or on benchmarking against simulated data. However, both approaches have important limitations and fail to capture the biological variability present in real datasets. FERRET is a framework for benchmarking single-cell GRN inference methods based on a simple biological assumption: independent estimates of the regulatory network from the same cellular state should resemble one another more closely than estimates from distinct cellular states. Rather than relying on incomplete or simulated ground truth, FERRET quantifies network robustness using two complementary metrics: *Robustness Area Under the Curve* (*RAUC*), an AUC-like measure of within-cell-type network similarity relative to between-cell-type similarity, and *Monotonicity*, which assesses the consistency of network similarity across edge-weight cutoffs. FERRET also supports biological validation through pathway enrichment analysis. We validate FERRET using experimentally derived ChIP-seq networks from B lymphocytes and fibroblasts as positive controls and randomly generated networks as negative controls, showing that biologically related networks receive high robustness scores whereas randomly generated networks receive scores consistent with chance. Finally, we apply FERRET to multiple GRN inference methods on real single-cell RNA-sequencing datasets to identify methods that produce the most robust, biologically informative regulatory networks.

**GRAPHICAL ABSTRACT:** 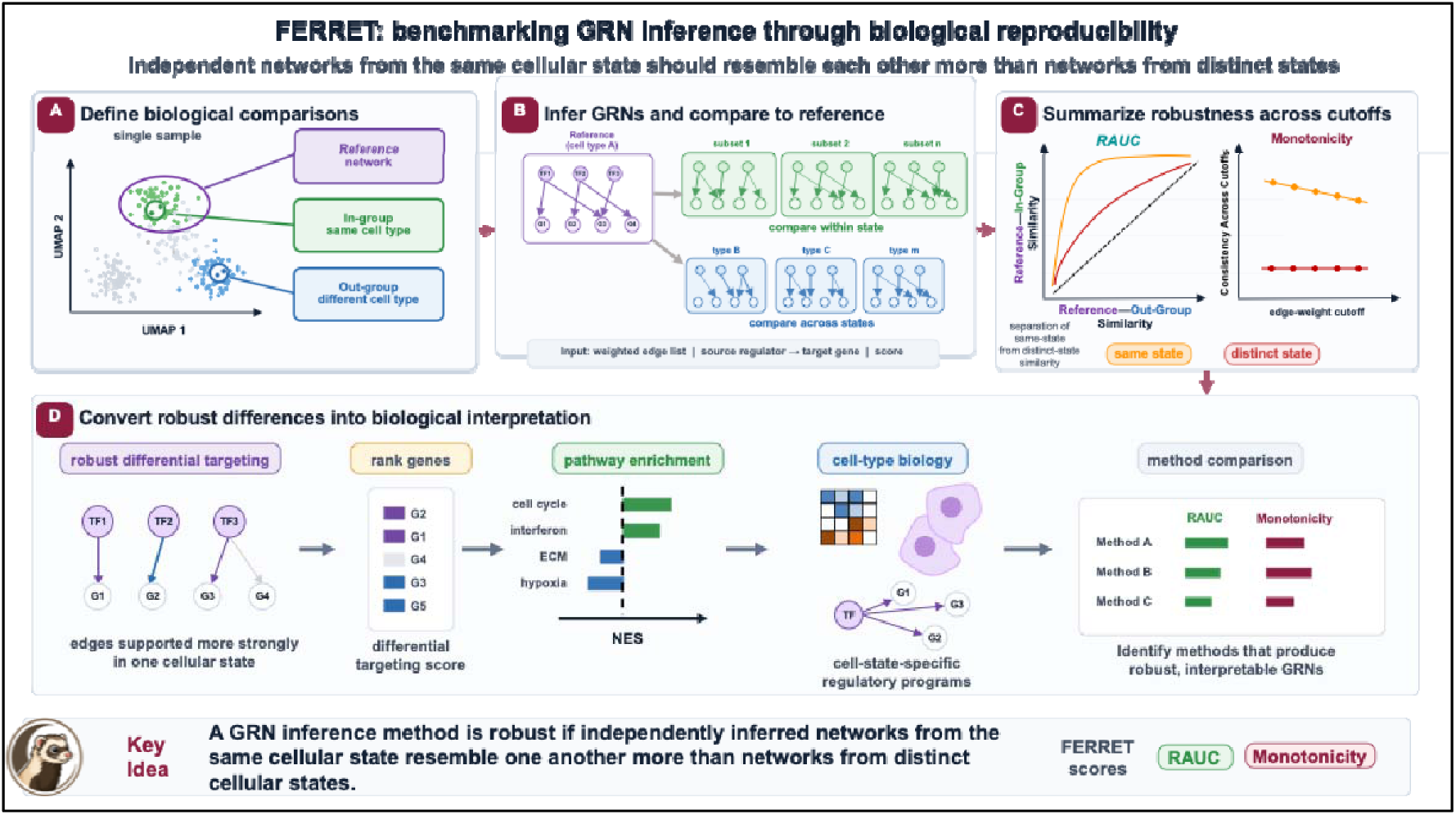

## INTRODUCTION

Gene regulatory networks (GRN) inference has become an essential approach for studying transcriptional regulation, but evaluating the quality of inferred networks remains a fundamental challenge. GRN analysis complements transcriptomic analyses such as differential expression by addressing not only which genes change, but also why they change as a consequence of transcription factor (TF) and other regulator activity (1). Experimentally constructing genome-wide GRNs would require measuring the binding or activity of every potential regulator across every relevant biological context, which is not currently feasible. Moreover, regulatory networks are expected to differ among cells, cell types, biological states, and over time. Consequently, statistical and computational methods are generally used to infer GRNs from gene expression and other molecular data (1–5). The advent of single-cell transcriptomics has led to a proliferation of methods for inferring cell-type-specific GRNs to identify the regulatory mechanisms underlying cellular heterogeneity. The rapid growth in GRN inference methods has created a corresponding need for rigorous and biologically meaningful approaches to evaluating their performance.

GRN inference methods differ considerably in how they represent regulatory relationships. Most model transcription as interactions between transcription factors (TFs) or other regulators and their target genes. Some explicitly distinguish regulators from targets, producing bipartite networks, while others allow genes to function as both regulators and targets, resulting in non-bipartite networks. Other approaches identify associations among genes without explicitly inferring regulator–target relationships masking potential causal associations. Some methods also incorporate prior biological knowledge, including TF binding motifs, protein–protein interactions, and curated regulatory networks. Examples of methods representing these different approaches are provided in Supplementary File 1. In this study, we exclude methods such as WGCNA (6,7), which infer gene co-expression networks rather than explicit regulator–target relationships. The diversity of modeling assumptions and network representations makes objective comparison of GRN inference methods particularly challenging.

Several analytical frameworks have been developed to evaluate the performance of GRN inference methods, each relying on different assumptions about what constitutes a correct regulatory network. The most widely used framework is BEELINE (Benchmarking Gene Regulatory Network Inference from Single-Cell Transcriptomic Data) (8), which benchmarks methods using ChIP-seq-derived networks, curated regulatory models, and simulated data, with metrics such as Area Under the Curve (AUC) and stability across repeated analyses. Other approaches rely more heavily on biological validation. For example, the developers of SCENIC (9) evaluated inferred GRNs based on expert interpretation and their ability to recover expected regulatory changes following transcription factor knockdown. Keyl *et al*. proposed a reproducibility-based evaluation for scGeneRAI by comparing similarities among networks inferred from the same individual with those inferred across individuals within a cell type (10).

While these approaches provide valuable measures of performance, each has important limitations. Simulated data may fail to capture the complexity of real biological systems and may favor methods whose assumptions resemble those used during simulation. Curated regulatory networks, ChIP-seq datasets, and perturbation experiments represent only a small fraction of regulatory interactions and are often generated in biological contexts distinct from those under study. Likewise, biological interpretation of inferred networks, while valuable, is inherently *a posteriori* and can bias evaluation toward previously characterized biology, potentially overlooking previously undescribed but biologically valid and relevant regulatory relationships (11). Finally, reproducibility across individuals reflects both methodological performance and inter-individual biological variation, making it difficult to isolate methodological robustness from the effects of biological heterogeneity.

To address these limitations, we developed FERRET (Framework to Evaluate Robustness in Regulatory Networks Using Heterogeneous Cell Types), a framework for evaluating GRN inference methods based on a simple biological assumption: independent estimates of the regulatory network from the same cellular state should resemble one another more closely than estimates from distinct cellular states (see Graphical Abstract). Rather than evaluating recovery of incomplete ground truth or relying on biological interpretation after inference, FERRET assesses whether GRN inference methods consistently recover biologically coherent regulatory structure by comparing similarities among networks inferred from different subsets of cells within the same cell type to similarities among networks inferred from different cell types from the same sample. Because these comparisons are performed within an individual sample, FERRET minimizes the influence of inter-individual variation while preserving biologically meaningful differences between cell types. FERRET summarizes these comparisons using two complementary metrics: the *Robustness Area Under the Curve* (*RAUC*), which quantifies robustness to within-cell-type biological variation, and *Monotonicity*, which measures the consistency of network similarity across edge-weight cutoffs; collectively, we refer to these as FERRET scores.

We evaluated FERRET using experimentally derived regulatory networks, simulated negative controls, and multiple cancer single-cell RNA-sequencing datasets. These analyses demonstrate that FERRET distinguishes biologically meaningful regulatory structure from random network organization and provides a complementary framework for benchmarking GRN inference methods based on biological reproducibility rather than recovery of incomplete ground truth.

### Overview of the FERRET framework

FERRET evaluates the robustness of gene regulatory network (GRN) inference methods by quantifying how well they preserve biologically meaningful relationships among inferred networks. Specifically, FERRET asks whether GRNs inferred from biologically related samples are consistently more similar to one another than to GRNs inferred from biologically distinct samples. Rather than evaluating a single edge-weight threshold, FERRET measures this separation across a range of edge-weight cutoffs using two complementary statistics: the *Robustness Area Under the Curve* (*RAUC*), which quantifies the overall separation between in-group and out-group networks, and the *Monotonicity* score, which measures the stability of that separation across thresholds.

FERRET operates on collections of GRNs inferred using the network inference method under evaluation. For each analysis, one or more networks are designated as reference GRNs and compared with biologically related in-group networks and biologically distinct out-group networks. GRNs are provided as weighted adjacency lists containing source node, target node, and edge-weight columns. The framework is agnostic to both the GRN inference method and the network similarity metric, allowing users to evaluate a broad range of inference approaches. Because different inference methods produce edge weights with different scales and distributions, FERRET evaluates robustness across user-defined edge-weight cutoffs rather than relying on a single threshold. Users may additionally specify whether negative edge weights represent weak evidence for a regulatory interaction or transcriptional inhibition and whether similarities are computed using raw or percentile-scaled edge weights. FERRET produces both the robustness curve and the corresponding *RAUC* and *Monotonicity* scores for each comparison.

#### Robustness Area Under the Curve (RAUC)

A robust GRN inference method should consistently place biologically related networks closer together than biologically distinct networks, regardless of the edge-weight threshold used to define the network. *RAUC* quantifies this behavior by summarizing the separation between in-group and out-group similarities across all edge-weight cutoffs.

For a given reference GRN, *g*_ref_, a set of in-group GRNs, *G*_in_, and a set of out-group GRNs, *G*_out_, for a single sample, s, and a user-defined set of edge-weight cutoffs, *C*, *RAUC* is calculated as

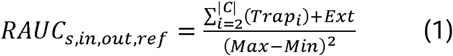

where *Trap_i_*, is the trapezoidal area between adjacent edge-weight cutoffs, *i* and *i* + 1, Ext is an extension term that accounts for cases in which the maximum in-group similarity exceeds the maximum out-group similarity, and Max and Min denote the overall maximum and minimum similarities observed across all in-group and out-group comparisons. Formal definitions of these quantities are provided in Supplementary Equation 1.

Equation (1) extends the conventional trapezoidal calculation of area under the curve by integrating over the observed range of network similarities rather than over the interval from 0 to 1. This normalization accommodates inference methods that produce different similarity distributions while preserving the biological interpretation of the resulting scores. Across multiple reference networks or cell types, *RAUC* scores may be consolidated by binning similarity values across edge-weight cutoffs, as described in Supplementary Equation 2.

Unlike receiver operating characteristic (ROC) or precision-recall analyses, FERRET evaluates relationships among complete inferred networks rather than individual regulatory interactions. Consequently, the similarity axes span the observed range of observed network similarities rather than the interval from 0 to 1. This formulation reflects the biological expectation that GRNs inferred from biologically related samples will share a substantial core of regulatory interactions while remaining neither identical because of biological heterogeneity nor completely dissimilar because essential cellular processes are broadly conserved (12,13).

Like conventional AUC statistics, *RAUC* ranges from 0 to 1. Values approaching 1 indicate that the reference GRN remains consistently more similar to biologically related networks than to biologically distinct networks across the full range of edge-weight cutoffs. A value near 0.5 indicates little or no separation between the two groups, whereas values below 0.5 indicate that the reference network is, on average, more similar to out-group than to in-group networks.

#### Monotonicity

*RAUC* summarizes the overall separation between in-group and out-group networks but does not describe how that separation changes as progressively weaker regulatory interactions are incorporated into the comparison. Two inference methods may therefore produce similar *RAUC* values while exhibiting markedly different behavior across edge-weight thresholds. We therefore introduce the *Monotonicity* score to quantify the stability of the robustness curve.

*Monotonicity* is calculated from the ordered in-group and out-group similarities that define the robustness curve. Let S_in_ and S_out_ denote the sorted in-group and out-group similarity values, respectively, as defined in Supplementary Equation 1. *Monotonicity* is calculated as,

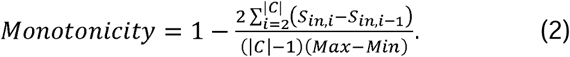

Monotonicity ranges from 0 to 1, with higher values indicating that in-group and out-group similarities change smoothly as edge-weight thresholds vary and lower values indicating abrupt changes in the robustness trajectory. Like *RAUC*, *Monotonicity* scores may be consolidated across multiple reference networks by binning similarity values across cutoffs.

### Joint Interpretation of *RAUC* and *Monotonicity*

*RAUC* and *Monotonicity* provide complementary measures of robustness (Figure 1). High values for both metrics indicate that biologically related networks remain well separated from biologically distinct networks and that this separation is maintained consistently across edge-weight thresholds. High *RAUC* together with lower *Monotonicity* indicates good overall separation but increased sensitivity to threshold selection, suggesting that the inferred edge weights are distributed inconsistently across thresholds. Conversely, lower *RAUC* with high *Monotonicity* indicates weaker overall biological separation that is nevertheless stable across thresholds. Low values for both metrics indicate poor separation, together with substantial sensitivity to threshold selection, reflecting limited robustness and instability. The joint interpretation of *RAUC* and *Monotonicity* is summarized in Table 1.

**Figure 1:**
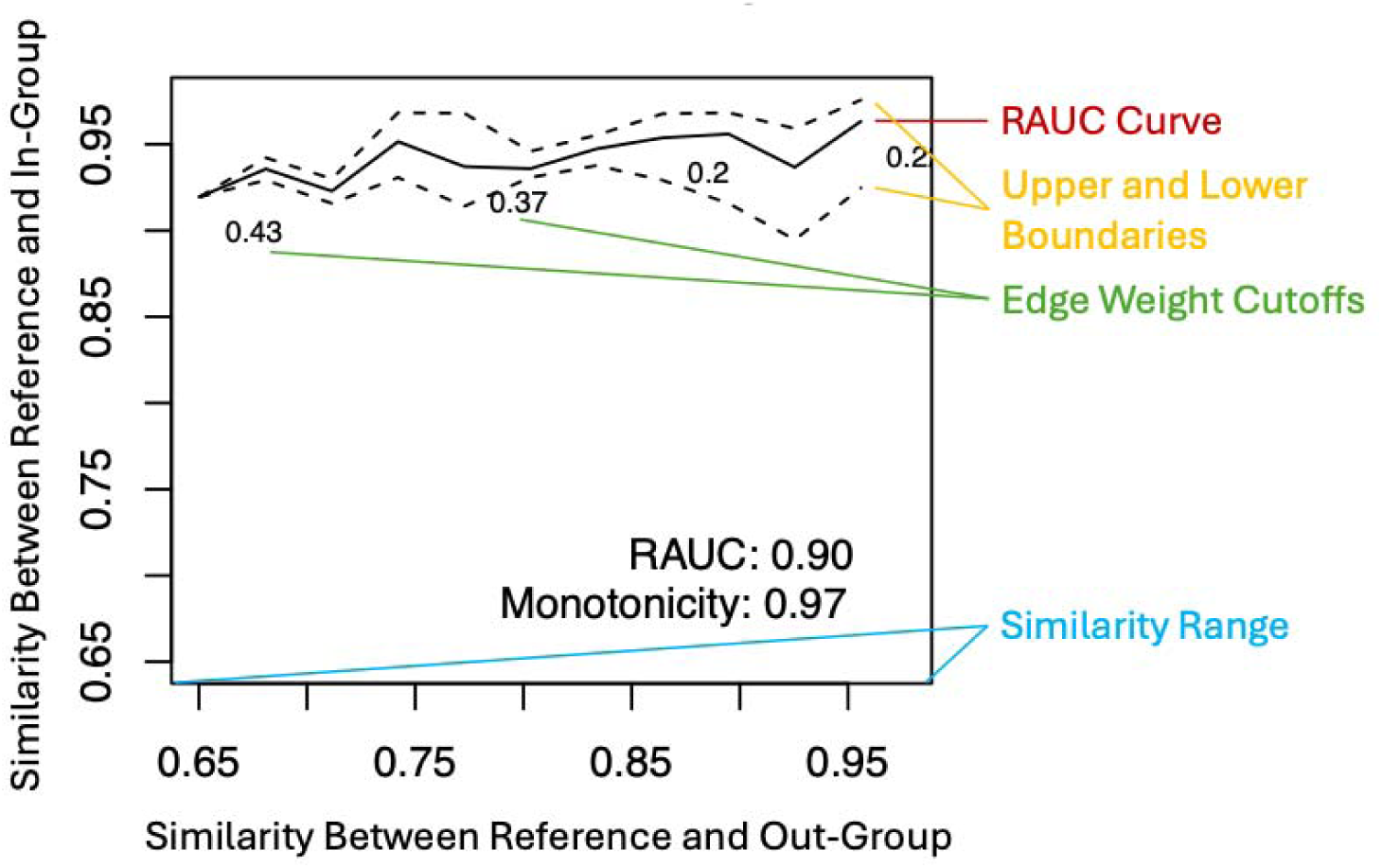
Example *RAUC* curve. As the cutoff for network edge weighs increases, the similarity between the reference and In-Group decreases slightly (in this case dropping from about 0.95 to 0.92) while the similarity between the reference and the Out-Group drops substantially (falling from 0.95 to 0.65). Overall *RAUC* and *Monotonicity* scores in the lower right.

**Table 1.** Joint interpretation of the *Robustness Area Under the Curve* (*RAUC*) and *Monotonicity* scores. RAUC quantifies the overall separation between biologically related (in-group) and biologically distinct (out-group) gene regulatory networks across edge-weight cutoffs, whereas Monotonicity measures the stability of that separation as progressively weaker regulatory interactions are incorporated into the analysis. Together, these complementary metrics distinguish methods that produce strong and stable biological separation from those whose performance is weaker or more sensitive to the choice of edge-weight cutoff.

|  | High *Monotonicity* | Low *Monotonicity* |
| --- | --- | --- |
| High *RAUC* | **Ideal performance.** Strong biological separation that is maintained consistently across edge-weight thresholds. | **Good but threshold-dependent.** Strong overall separation, but robustness depends on the selected cutoff. |
| Low *RAUC* | **Consistent but weak.** Stable behavior across thresholds, but limited ability to distinguish biologically related from unrelated networks. | **Poor performance.** Weak biological separation together with instability across thresholds. |

## MATERIALS AND METHODS

### Overview of the Benchmarking Strategy

We evaluated FERRET using three complementary benchmarking strategies designed to assess its ability to distinguish biologically meaningful differences in gene regulatory networks (GRNs) from random variation and to compare the robustness of existing GRN inference methods. First, we generated biologically meaningful positive controls using regulatory networks constructed from publicly available ChIP-seq data collected in fibroblasts and B-lymphocytes. Because these cell types exhibit distinct regulatory programs, we expected networks inferred from the same cell type to be consistently more similar than networks inferred from different cell types, resulting in high RAUC scores. Second, we generated negative controls using randomly simulated directed networks for which no biologically meaningful grouping exists, and, therefore, *RAUC* scores near 0.5 are expected. Together, these analyses established the expected behavior of FERRET under biologically meaningful and random conditions.

We next used FERRET to benchmark multiple GRN inference methods using three independent single-cell RNA sequencing (scRNA-seq) datasets representing glioblastoma (GBM), renal cell carcinoma (RCC), and small cell lung cancer (SCLC). For each dataset, we inferred GRNs from biologically defined cell populations, constructed reference, in-group, and out-group network sets, and evaluated their robustness using the *RAUC* and *Monotonicity* statistics. Because FERRET is independent of the underlying network inference algorithm, this benchmarking strategy enabled direct comparison of methods that differ substantially in their underlying assumptions, modeling approaches, and edge-weight distributions.

Finally, we evaluated the biological plausibility of the networks identified by each inference method. Differential regulatory interactions identified by FERRET were used to generate differential targeting scores, which were subsequently analyzed for pathway enrichment. This analysis provided a complementary biological assessment of each inference method by determining whether the inferred regulatory differences recapitulated known functional distinctions among cell types.

### Positive and Negative Control Generation

#### Positive Control Generation

To establish the expected behavior of FERRET under biologically meaningful conditions, we generated positive control gene regulatory networks (GRNs) from publicly available chromatin immunoprecipitation sequencing (ChIP-seq) experiments performed in fibroblasts (NHDF, NHLF, and IMR90 cell lines) and B-lymphocytes (isolated CD20+ cells and the GM12878 cell line). These cell types possess distinct regulatory programs and therefore provide an appropriate benchmark in which networks derived from the same cell type are expected to be consistently more similar than networks derived from different cell types.

ChIP-seq datasets were downloaded from the Cistrome Data Browser (Cistrome DB) (14). Samples were required to pass all Cistrome quality-control criteria, be aligned to either the hg19 or hg38 GENCODE reference genome (15), and be provided in Browser Extensible Data (BED), BROADPEAK, or NARROWPEAK format with standard peak scores ranging from 1 to 1000 (16). To facilitate direct comparison between cell types, we retained only transcription factors assayed in both fibroblasts and B-lymphocytes. After filtering, four fibroblast samples and six B-lymphocyte samples remained, representing the transcription factors *H2AZ*, *POLR2A*, and *RAD21* (Supplementary Table 1).

Using BEDTools (17), we retained only peaks located within promoter regions of protein-coding genes, defined as 750 bp upstream to 250 bp downstream of each transcription start site. Y chromosome genes and mitochondrial genes were excluded from the analysis (Supplementary Table 2). For each cell type, the reference network (*g*_ref_) was constructed by combining all available samples for each transcription factor. An edge was included if the corresponding TF-gene interaction was present in any sample. Peak scores for duplicate TF-gene pairs were first consolidated within each sample by retaining the maximum peak score and were then averaged across all samples to generate a single edge weight.

To construct the in-group network set (*G*_in_) for fibroblasts, we performed leave-one-out reconstruction of the *H2AZ* fibroblast samples while retaining the single *POLR2A* and *RAD21* samples in every network. The out-group network set (*G*_out_) for fibroblasts was generated by performing leave-one-out reconstruction across all *H2AZ*, *POLR2A*, and *RAD21* B-lymphocyte samples. For each analysis, the fibroblast *G*_in_networks served as the B-lymphocyte *G*_out_ networks and vice versa.

#### Negative Control Generation

To characterize FERRET under conditions in which no biologically meaningful network structure exists, we generated negative controls using simulated random networks. One hundred directed Barabási-Albert networks (18), each containing 1,000 nodes and 1,000 edges, were generated using the *igraph* R package (19). Edge weights were assigned independently by sampling from a uniform distribution on the interval [0.1, 1.0].

The simulated networks were partitioned into five folds. Within each fold, one network was designated as the reference network (*g*_ref_), while the remaining networks were assigned randomly to the in-group (*G*_in_) and out-group (*G*_out_). Because the assignment was random, no systematic differences between the two groups were expected, providing a negative control for which RAUC values should be centered near 0.5.

The simulated networks were directed but not bipartite. Because many GRN inference methods generate bipartite transcription factor-target networks, we transformed the simulated networks into equivalent bipartite representations using a one-to-one mapping procedure that we term bipartitification (defined in Supplementary File 1). We show in Supplementary File 1 that this transformation preserves the Jaccard, In-Degree, and Out-Degree similarity measures used by FERRET (Theorems 1-3 in Supplementary File 1) while producing valid bipartite networks. Consequently, the results obtained from the simulated directed networks generalize directly to bipartite regulatory networks.

### Expression Datasets

We analyzed three independent single-cell RNA sequencing (scRNA-seq) benchmark datasets representing distinct tumor types and biological contexts: glioblastoma (GBM), renal cell carcinoma (RCC), and small cell lung cancer (SCLC). The GBM and RCC datasets were obtained from the Clinical Proteomic Tumor Analysis Consortium (CPTAC) (20), whereas the SCLC dataset was obtained from the Human Tumor Atlas Network (HTAN) (21).

For the CPTAC datasets, biospecimen information, clinical annotations, sample metadata, and sample sheets were downloaded from the Genomic Data Commons (GDC) Data Portal (22), and the corresponding scRNA-seq data were retrieved using the GDC Data Transfer Tool (23). The HTAN SCLC scRNA-seq dataset was downloaded directly from the HTAN data portal.

### Expression Data Set Preprocessing

Single-cell RNA sequencing (scRNA-seq) datasets were processed using the Bioconductor *SingleCellExperiment* framework (24). Data were imported using the *SingleCellExperiment* and *loomR* R packages (25), annotated with gene symbols and genomic locations using *scater* (26), and normalized by estimating cell-specific size factors with *scran* (27). Quality control metrics were computed using *scater*.

To improve data quality and ensure consistent gene annotation across datasets, duplicated gene symbols were resolved by retaining the transcript with the greatest expression for each annotated gene. Cells with fewer than 1,000 detected genes or with at least 50% mitochondrial reads were excluded from subsequent analyses.

Cell types were assigned using the *scMRMA* R package (28) with *PanglaoDB* as the reference database (29). We retained neurons, interneurons, oligodendrocytes, and macrophages for the CPTAC GBM dataset; podocytes and macrophages for the CPTAC RCC dataset; and macrophages, endothelial cells, fibroblasts, natural killer (NK) cells, T cells, and B cells for the HTAN SCLC dataset. These cell populations were selected because they were consistently represented across samples and provided sufficient cell numbers for downstream pseudotime estimation and GRN inference.

### Pseudotime Trajectory Assignment

Pseudotime trajectories were inferred independently for each sample and cell type using the *Monocle3* R package (30). Cells were clustered, embedded in a low-dimensional manifold, and ordered along an inferred developmental trajectory. The resulting trajectories were projected onto Uniform Manifold Approximation and Projection (UMAP) embeddings (31) and evaluated using the expression patterns of established cell-type-specific marker genes.

Trajectory orientation was determined manually by identifying start and end points that were consistent with the expected progression of cell-type-specific marker gene expression. Cell-type-specific marker genes were identified as described in Supplementary File 1. We retained only samples for which biologically interpretable pseudotime trajectories could be defined for at least two cell types, thereby ensuring that downstream comparisons of GRN robustness reflected meaningful differences in regulatory dynamics rather than artifacts of trajectory inference.

### Dimensionality Reduction

Each scRNA-seq dataset was projected onto its first 100 principal components using the *irlba* R package to reduce dimensionality while preserving the dominant sources of transcriptional variation. GRN inference was subsequently performed using these low-dimensional representations rather than the full gene-expression matrices.

Benchmarking was standardized by randomly selecting 63 cells from each sample-cell type combination, corresponding to the smallest cell population that met the pseudotime quality-control criteria. For each sample and cell type, the resulting input to each GRN inference method consisted of a 100 × 63 matrix of principal component scores together with a corresponding vector of pseudotime labels. Standardizing both the dimensionality of the input data and the number of cells ensured that differences in computational performance and inferred network structure reflected differences among GRN inference methods rather than variation in input.

### Reference, In-Group, and Out-Group Designation

Reference, in-group, and out-group GRNs were constructed independently for each sample and pair of cell types. For each comparison, the reference network (*g*_ref_) was inferred from the full 100 × 63 reference matrix for one cell type. The in-group network set (*G*_in_) consisted of GRNs inferred from resampled subsets of the same cell type, whereas the out-group network set ( *G*_out_) consisted of GRNs inferred from resampled subsets of the comparison cell type. This design ensured that in-group comparisons measured within-cell-type variability, while out-group comparisons quantified differences between biologically distinct cell types.

For the CPTAC datasets, resampling was performed using ten folds with five random splits per fold. Each split contained 57 of the 63 cells from the corresponding sample-cell type combination, resulting in 50 in-group and 50 out-group networks for every reference network.

For the HTAN dataset, resampling was performed using five folds with a single split per fold. Each reference network was therefore compared with five in-group and five out-group networks.

This resampling strategy generated multiple independent realizations of each biological condition while maintaining a consistent reference network, allowing FERRET to quantify the robustness of GRN inference methods in the presence of biological and sampling variability.

### Metrics for Network Similarity

FERRET quantifies similarity between gene regulatory networks using three complementary metrics: Jaccard Similarity, In-Degree Similarity, and Out-Degree Similarity, defined in Supplementary Equations (3–5), respectively. Jaccard Similarity measures the overlap in regulatory interactions between two networks. In contrast, In-Degree Similarity compares the overall regulatory input received by each target gene, whereas Out-Degree Similarity compares the overall regulatory output of each source gene.

These metrics capture complementary aspects of network organization. Two networks may exhibit substantial overlap in individual regulatory interactions while differing in the extent to which specific genes are regulated or function as regulators, leading to distinct biological outcomes (1). Conversely, two networks may exhibit similar overall regulatory activity despite differing in the specific regulatory interactions that give rise to that activity. Evaluating all three similarity measures provides a more comprehensive assessment of network similarity than any single metric alone. It is worth noting that because FERRET is independent of the network similarity metric, additional similarity measures can be incorporated within the method’s benchmarking framework.

### Pathway Enrichment Analysis and Interpretation

FERRET identifies regulatory differences between cell types by first constructing consensus networks for the in-group and out-group comparisons. For each GRN inference method, the common regulatory networks shared among the in-group, n (*G*_in_), and out-group, n (*G*_out_), networks were identified for each sample and cell type as defined in Supplementary Equations (6) and (7). A differential regulatory network was then constructed by comparing the two consensus networks (Supplementary Equation 8), with an extension for dimensionally reduced networks described in Supplementary Equation (9). Differential targeting scores were subsequently computed for every gene using Supplementary Equation (10).

For dimensionally reduced networks, such as those inferred from principal component representations of the CPTAC datasets, gene-level edge weights were reconstructed by multiplying the edge weights in the reduced network by the corresponding source and target loading matrices, as described in Supplementary File 1.

Pathway enrichment analysis was performed on the resulting differential targeting scores using the *fgsea* R package (32). Gene sets were provided by the user and included pathway collections such as the Molecular Signatures Database (MSigDB) (33). Genes were ranked according to their differential targeting scores prior to enrichment analysis.

Significantly enriched pathways were first grouped into functional categories. The biological relevance of each category was then evaluated by assessing its concordance with established cell-type biology using a structured literature review described in Supplementary File 1. This analysis provides an independent biological assessment of each GRN inference method by determining whether the regulatory differences identified by FERRET recapitulate known functional distinctions between cell types.

### Gene Regulatory Network Inference Methods

Candidate GRN inference methods were screened using the inclusion criteria summarized in Table 2. These criteria were designed to ensure meaningful and reproducible comparisons across diverse inference paradigms while accounting for the different characteristics of dimensionally reduced and genome-wide expression data.

**Table 2.** Inclusion criteria for gene regulatory network (GRN) inference methods included in the FERRET benchmark.

| **Criterion** | **Dimensionally Reduced Expression Data** | **Genome-wide Expression Data** |
| --- | --- | --- |
| **Ground truth regulatory network** | Not required | Not required |
| **Publicly available software implementation** | Required | Required |
| **Weighted network output** | Required (confidence or interaction strength reported for every inferred edge) | Required (confidence or interaction strength reported for every inferred edge) |
| **Applicable to scRNA-seq data** | Required | Required |
| **Applicable to single-timepoint data** | Required | Required |
| **External gene-level regulatory information** | Not permitted | Permitted |
| **Primary selection criterion** | Compatibility with reduced-dimensional inputs | Completion within 48 hours on a representative genome-wide dataset |

Six methods—GRISLI (34), LEAP (35), PIDC (36), scGeneRai (10), scSGL (37), and SINGE (38)—were evaluated using dimensionally reduced expression data from the CPTAC glioblastoma (GBM) and renal cell carcinoma (RCC) datasets. These analyses were performed on matrices consisting of the first 100 principal components for 63 cells per sample-cell type combination. These methods were selected because they do not require gene-level regulatory priors and either do not scale efficiently to genome-wide analyses or were designed for more modest-sized inputs.

SCORPION (39) and SCENIC (40) were evaluated using genome-wide expression data from the HTAN small cell lung cancer (SCLC) dataset. Both methods rely on gene-level regulatory information, making them incompatible with the reduced-dimensional representation. They were retained for the genome-wide benchmark because they completed inference on a representative genome-wide macrophage dataset within the required 48-hour runtime threshold. The remaining six methods exceeded this runtime threshold on genome-wide data, and GRISLI could not be evaluated because its estimated array-size requirement was approximately 31,610 GB.

Overall, the benchmark design was determined by both methodological compatibility and computational scalability. SCORPION and SCENIC require gene-level resolution and were sufficiently efficient for genome-wide analyses, whereas GRISLI, LEAP, PIDC, scGeneRai, scSGL, and SINGE required dimensionality reduction to enable practical benchmarking.

All benchmarking analyses were performed on an Amazon Elastic Compute Cloud (EC2) instance provided by Amazon Web Services (AWS). The computational environment consisted of an hpc6id.32xlarge instance with 64 vCPUs, 1,204 GB of memory, and 200 Gbps network bandwidth.

## RESULTS

FERRET evaluates the robustness of gene regulatory network (GRN) inference methods using two complementary metrics, *Robustness Area Under the Curve (RAUC)* and *Monotonicity*. We first evaluated whether these metrics behaved as expected using positive and negative controls with known biological properties. We then benchmarked existing GRN inference methods on dimensionally reduced and genome-wide single-cell RNA sequencing datasets to determine whether FERRET distinguishes methodological differences and identifies biologically meaningful regulatory programs.

### FERRET Behaves as Expected in Positive and Negative Controls

#### Positive Controls

We evaluated FERRET using experimentally derived ChIP-seq regulatory networks from fibroblasts and B-lymphocytes, two cell types with well-established differences in regulatory biology. For each cell type, the reference network, *g*_ref_, was constructed from all available samples, whereas the in-group, *G*_in_, and out-group, *G*_out_, networks were generated using leave-one-out reconstructions from the same and opposing cell types, respectively. Because regulatory networks derived from the same cell type are expected to be substantially more similar than those derived from different cell types, we expected both *RAUC* and *Monotonicity* to be high.

FERRET behaved as expected across all similarity metrics (Figure 2). *RAUC* scores ranged from 0.85 to 0.96, whereas *Monotonicity* ranged from 0.95 to 1.00, indicating that regulatory networks inferred from the same cell type remained consistently more similar across all edge-weight cutoffs than networks inferred from different cell types.

**Figure 2:**
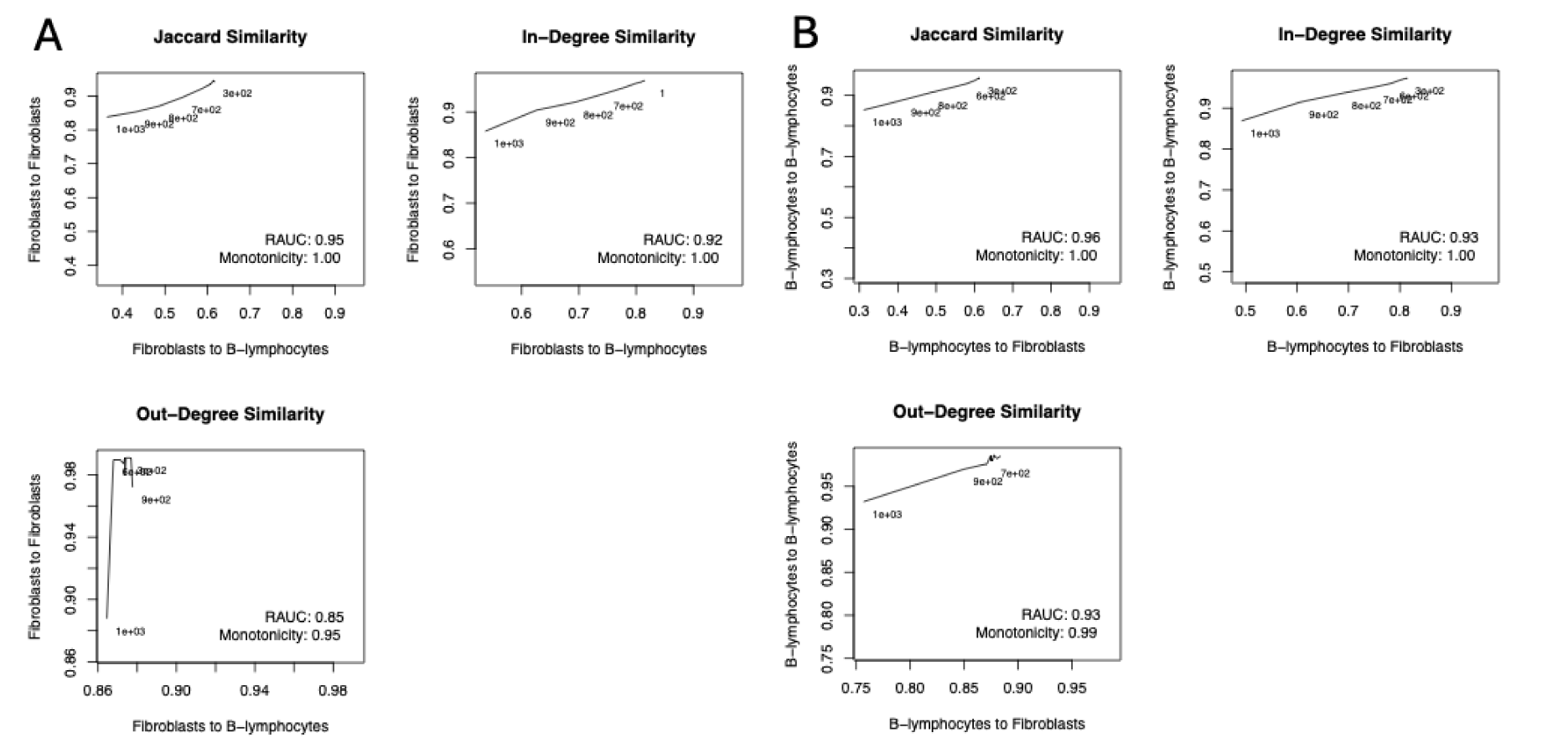
RAUC curves for ChIP-seq data when evaluating (A) fibroblasts against B-lymphocytes and (B) B-lymphocytes against fibroblasts. For all similarity metrics, *RAUC* is at least 0.85 and *Monotonicity* is at least 0.95. Moreover, as score cutoffs become more stringent, overall similarity decreases.

Differential targeting analysis further demonstrated that these robustness scores reflected biologically meaningful differences rather than purely topological distinctions (Supplementary Tables 3–4). Thirty-two pathways were more strongly targeted in B-lymphocytes than fibroblasts, including cytokine signaling in immune system, B cell receptor signaling, signaling by interleukins, leukocyte transendothelial migration, and multiple pathways associated with B-cell activation and immune function. Conversely, 138 pathways exhibited greater targeting in fibroblasts, including degradation of the extracellular matrix, collagen degradation, collagen formation, laminin interactions, cell junction organization, and additional pathways involved in extracellular matrix organization and tissue remodeling. Together, these results demonstrate that high *RAUC* and *Monotonicity* scores correspond to biologically meaningful differences between cell types.

#### Negative Controls

We next evaluated FERRET using randomly generated networks with random assignments of reference, in-group, and out-group networks. Under these conditions, the reference network is expected to be equally similar to the in-group and out-group networks, producing *RAUC* values near 0.5. Because increasing edge-weight cutoffs progressively sparsify all random networks, both in-group and out-group similarities are expected to decrease consistently across cutoffs, resulting in high Monotonicity scores.

FERRET again behaved as expected (Figure 3). *RAUC* scores remained centered near 0.5 with minimal variation across random assignments, while Monotonicity remained uniformly high, reflecting the consistent decline in similarity as networks became progressively sparser. These results demonstrate that FERRET distinguishes biologically meaningful regulatory structure from randomly generated networks while producing the expected behavior under both positive and negative control conditions.

**Figure 3:**
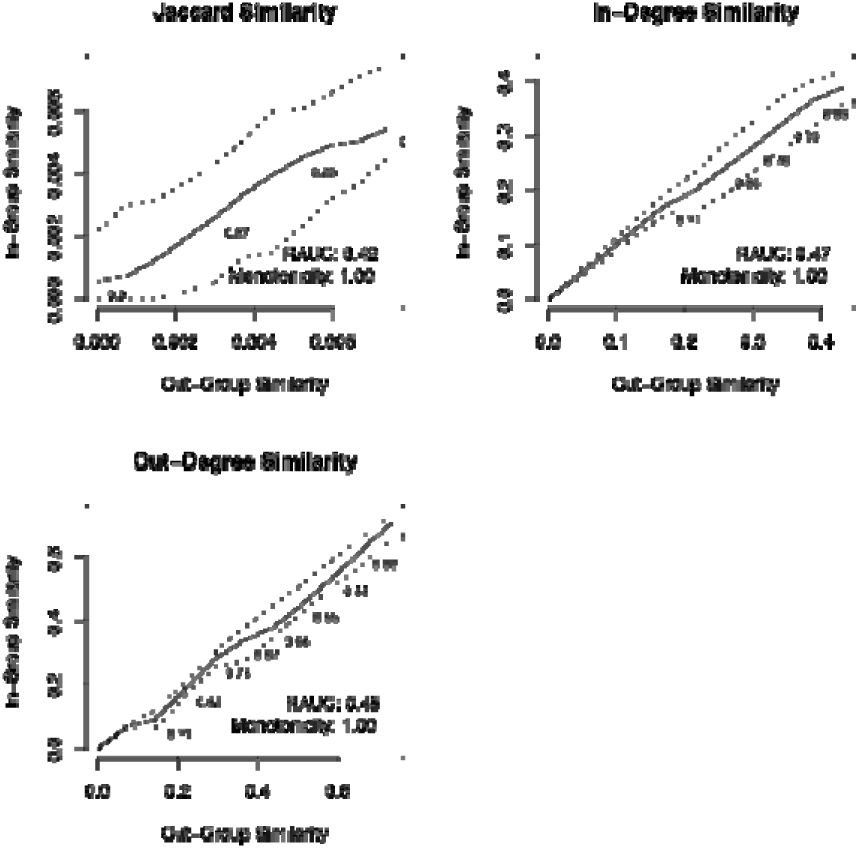
RAUC curves for random networks. For all similarity metrics, random networks assigned to random, , and groupings achieved *RAUC* scores between 0.4 and 0.5, consistent with the expected values for random performance in AUC scores. Cutoffs are labelled in smaller font along the curve.

### FERRET Distinguishes Methodological Strengths and Weaknesses of GRN Inference Methods

We next applied FERRET to benchmark six GRN inference methods—GRISLI, LEAP, PIDC, scGeneRai, scSGL, and SINGE—using the dimensionally reduced CPTAC datasets (Figure 4). Because these methods differ substantially in their underlying statistical formulations, we expected FERRET to distinguish differences in robustness and stability across similarity metrics and cell types.

**Figure 4.**
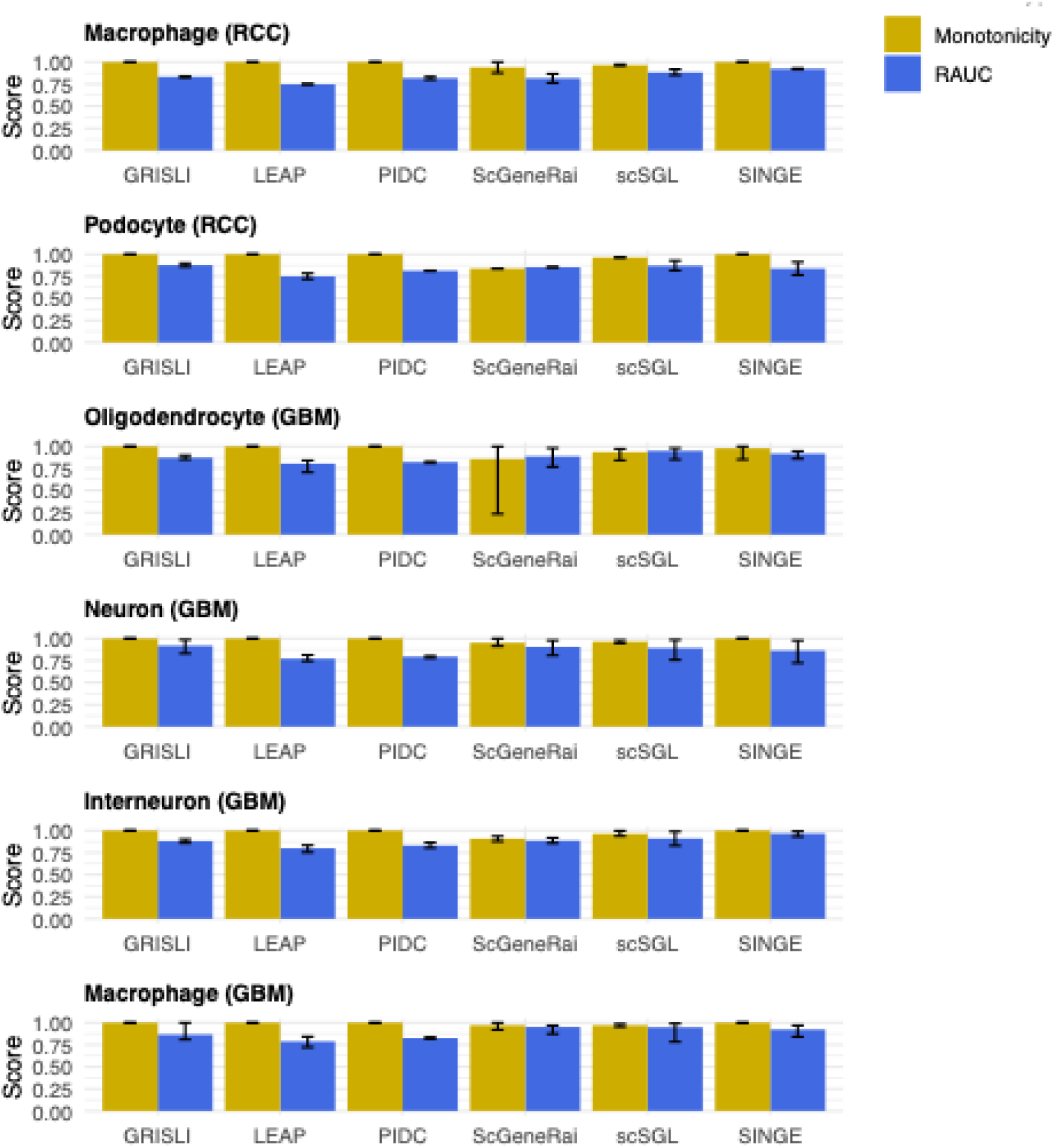
RAUC and Monotonicity results for all methods evaluated on dimensionality reduced CPTAC data using Jaccard similarity.

#### Robustness

scGeneRai, GRISLI, and scSGL achieved the highest *RAUC* scores using the Jaccard and Out-Degree similarity metrics. GRISLI performed less favorably when evaluated using the In-Degree metric, indicating that although it consistently recovered cell-type-specific regulatory interactions and transcription factor regulatory profiles, it was less effective at distinguishing differences in gene-targeting profiles. In contrast, SINGE consistently achieved the lowest RAUC scores using the Out-Degree metric but performed substantially better using the In-Degree metric, suggesting that it more effectively captures differences in target-gene regulation than transcription factor activity in dimensionally reduced space (Figure 4; Supplementary Figures 1-2; Supplementary Table 5).

These differences are consistent with the underlying methodological design of each algorithm. scGeneRai, scSGL, and GRISLI employ nonlinear multivariate models capable of capturing complex relationships among multiple regulators, whereas PIDC and LEAP primarily infer pairwise regulatory relationships using information-theoretic or correlation-based approaches. The improved robustness of the nonlinear methods suggests that modeling higher-order regulatory interactions provides an advantage when distinguishing cell-type-specific regulatory structure. GRISLI’s comparatively weaker performance using the In-Degree metric is also consistent with its sparsity-promoting formulation, which favors a relatively small set of regulators for each target gene and consequently produces more similar target-gene regulatory profiles across cell types.

#### Stability Across Edge-Weight Cutoffs

PIDC consistently achieved the highest Monotonicity scores across similarity metrics and cell types, indicating that its relative ranking of in-group and out-group similarities remained highly stable across edge-weight cutoffs. In contrast, scGeneRai exhibited substantially lower Monotonicity, particularly for oligodendrocytes using the Jaccard and In-Degree metrics. Because *Monotonicity* reflects the consistency of edge-weight distributions across repeated network inference, these differences likely arise from the underlying scoring schemes used by each method. PIDC computes edge weights from information gain, producing values on a common numerical scale across networks, whereas scGeneRai derives edge weights from neural network model coefficients, whose magnitudes are less directly comparable across independently trained models.

#### Comparison with BEELINE

FERRET rankings showed broad agreement with curated-network benchmarks reported by BEELINE. Using curated regulatory networks, BEELINE ranked PIDC highest overall, followed by GRISLI, SINGE, and LEAP. FERRET produced a similar ordering using the In-Degree similarity metric, differing primarily in the stronger performance of SINGE relative to PIDC. Rankings based on the Out-Degree metric also broadly agreed with BEELINE, with the notable exception of PIDC.

These differences reflect the distinct goals of the two benchmarking frameworks. BEELINE evaluates the recovery of known regulatory interactions using curated reference networks, whereas FERRET evaluates the robustness of methods in distinguishing biologically meaningful regulatory differences without requiring a curated gold standard. Consequently, methods such as SINGE may identify biologically relevant regulatory changes that are absent from existing reference networks, leading to improved FERRET performance despite more modest agreement with curated interactions.

Together, these results demonstrate that FERRET not only recapitulates conclusions obtained from curated-network benchmarks but also provides complementary information by evaluating the robustness and biological relevance of inferred regulatory networks without requiring a gold-standard reference.

### Computational Scalability Limits Genome-wide Benchmarking

Genome-wide benchmarking requires GRN inference methods that are both computationally tractable and compatible with gene-level regulatory information. Consequently, we evaluated the computational performance of all candidate methods using macrophage scRNA-seq from a representative HTAN sample (RU682) and required completion within a 48-hour runtime threshold.

Only SCORPION and SCENIC completed inference within the allotted time (Table 3). SCORPION completed inference in 7,940 seconds (2.2 hours), whereas SCENIC required 25,673 seconds (7.1 hours). LEAP, PIDC, scGeneRai, scSGL, and SINGE all exceeded the 48-hour runtime threshold. GRISLI could not be evaluated because its estimated memory requirement exceeded 31 TB, making execution infeasible on the available computational infrastructure.

**Table 3.** GRN inference method runtimes tested on macrophages from a single sample. Note that GRISLI could not be evaluated for this sample because it required an array size of dimensionality 18,803 x 18,803 x 8 x 1,500, or approximately 31,610.1 GB.

| GRN Inference Method | Runtime |
| --- | --- |
| SCORPION | 7,940 s (2.2 hours) |
| SCENIC | 25,673 s (7.1 hours) |
| LEAP | > 48 hours |
| scSGL | > 48 hours |
| scGeneRai | > 48 hours |
| PIDC | > 48 hours |
| SINGE | > 48 hours |
| GRISLI | N/A |

Because both SCORPION and SCENIC meet the computational requirements for genome-wide analysis and operate directly on gene-level regulatory information, these two methods were used to infer GRNs for all HTAN samples and cell types. FERRET was then used to calculate *RAUC* and *Monotonicity* scores for each method and to perform pathway enrichment analyses based on differential targeting scores.

These results demonstrate that computational scalability remains a practical limitation for genome-wide GRN inference. Consequently, the subsequent genome-wide benchmark compares methods that are both computationally feasible and methodologically compatible with gene-level regulatory analyses.

### FERRET Differentiates SCORPION and SCENIC in Genome-wide GRN Inference

Following the computational scalability analysis, we evaluated SCORPION and SCENIC using genome-wide HTAN single-cell RNA sequencing data representing six major cell types. FERRET was used to compare the robustness of the inferred gene regulatory networks and to assess whether differences in network structure corresponded to biologically meaningful differences in regulatory targeting (Figure 5; Supplementary Figures 3–4; Supplementary Tables 6-7).

**Figure 5.**
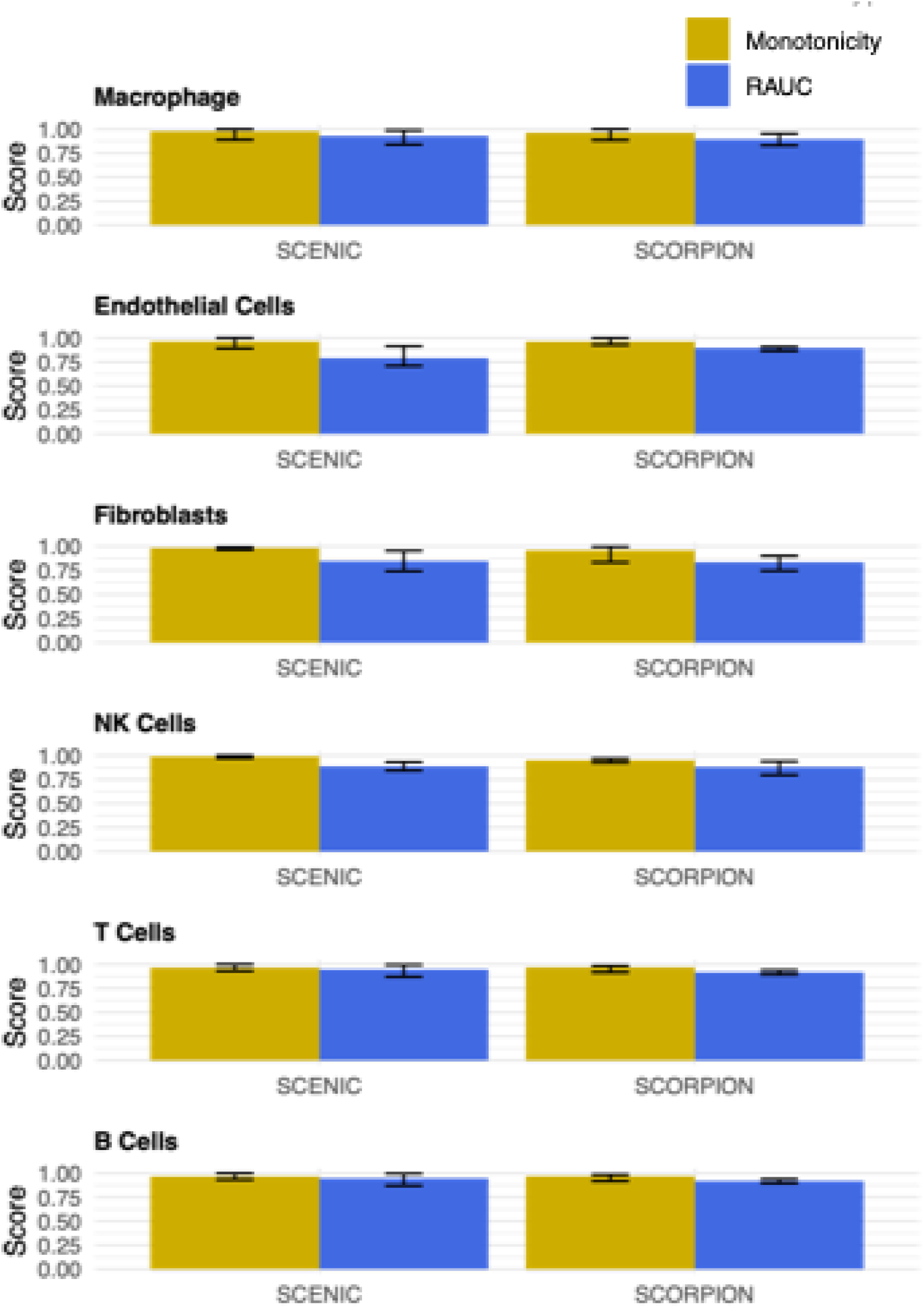
*RAUC* and *Monotonicity* results for all methods evaluated on genome-wide HTAN data using Jaccard similarity.

#### Robustness of Genome-wide GRN Inference

SCORPION and SCENIC exhibited similar RAUC and Monotonicity scores across cell types when evaluated using the Jaccard and In-Degree similarity metrics (Figure 5; Supplementary Figures 3–4; Supplementary Table 6). In contrast, SCENIC consistently achieved higher RAUC scores using the Out-Degree similarity metric, indicating improved discrimination of cell-type-specific transcription factor regulatory activity. Monotonicity remained high for both methods across all similarity metrics, suggesting that both methods assign edge weights consistently across edge-weight cutoffs despite differences in their underlying inference strategies.

Overall, these results indicate that SCORPION and SCENIC generate similarly robust genome-wide regulatory networks but differ in the aspects of regulatory biology that they capture. Whereas both methods distinguish cell-type-specific regulatory interactions, SCENIC more effectively captures differences in transcription factor activity across cell types.

#### Biological Interpretation of Differential Targeting

Differential targeting analysis followed by pathway enrichment identified substantially more enriched pathways in SCENIC-derived networks than in SCORPION-derived networks (Supplementary Table 7). To evaluate the biological relevance of these results, enriched pathways were grouped into functional categories and compared with published literature describing the biology of each cell type.

SCORPION identified a higher proportion of literature-supported pathway categories in fibroblasts and T cells, whereas SCENIC identified a higher proportion in B cells, endothelial cells, macrophages, and natural killer (NK) cells (Table 4). Although SCENIC generally identified a larger number of differentially targeted pathways, the proportion supported by the literature varied by cell type, suggesting that the two methods emphasize different aspects of regulatory biology rather than one method consistently outperforming the other.

**Table 4.** Literature Support for Enriched Pathways by Cell Type and GRN Inference Method in HTAN.

| Method | Cell Type (In-Group) | Number of Pathway Categories with Literature Support | Number of Pathway Categories without Literature Support | Percentage with Literature Support |
| --- | --- | --- | --- | --- |
| **SCENIC** | B Cells | 5 | 4 | 55.56% |
|  | Endothelial Cells | 1 | 10 | 9.09% |
|  | Fibroblasts | 27 | 31 | 46.55% |
|  | Macrophages | 24 | 29 | 45.28% |
|  | NK Cells | 10 | 26 | 27.78% |
|  | T Cells | 44 | 75 | 36.97% |
| **SCORPION** | B Cells | 1 | 5 | 16.67% |
|  | Endothelial Cells | 0 | 3 | 0.00% |
|  | Fibroblasts | 2 | 0 | 100.00% |
|  | Macrophages | 0 | 1 | 0.00% |
|  | NK Cells | 1 | 4 | 20.00% |
|  | T Cells | 2 | 3 | 40.00% |

These differences are consistent with the methodological design of the two algorithms. SCORPION incorporates transcription factor binding motifs as prior biological knowledge, constraining inferred regulatory interactions to relationships supported by known DNA binding. Consequently, differences between cell types arise primarily through changes in the strengths of established regulatory interactions. In contrast, SCENIC infers regulatory programs directly from patterns of gene co-expression and regulon activity, allowing greater flexibility in identifying context-dependent regulatory differences that may extend beyond currently curated transcription factor binding relationships.

Taken together, these results demonstrate that SCORPION and SCENIC provide complementary views of genome-wide gene regulation. SCORPION emphasizes conservative, prior-supported regulatory interactions, whereas SCENIC more readily identifies context-specific changes in transcription factor activity and downstream regulatory targeting. FERRET distinguishes these complementary methodological characteristics while simultaneously evaluating the robustness and biological relevance of inferred regulatory networks without requiring prior knowledge of the true underlying regulatory network.

## DISCUSSION

FERRET provides a framework for evaluating the robustness and biological relevance of gene regulatory network (GRN) inference methods without requiring prior knowledge of the underlying regulatory network. Rather than assessing recovery of a predefined set of regulatory interactions, FERRET evaluates whether inferred networks consistently preserve expected similarities within biologically related groups while distinguishing biologically distinct groups across a range of edge-weight cutoffs. By coupling the *Robustness Area Under the Curve (RAUC)* and *Monotonicity* metrics with differential targeting and pathway enrichment analyses, FERRET simultaneously evaluates both the reproducibility of inferred regulatory networks and the biological significance of the differences they identify.

The positive and negative control analyses demonstrate that FERRET behaves as expected under conditions where the anticipated outcome is known. Experimentally derived ChIP-seq networks from fibroblasts and B-lymphocytes produced high *RAUC* and *Monotonicity* scores together with pathway enrichments consistent with established cell-type biology. Conversely, randomly generated networks yielded *RAUC* scores near 0.5 and consistently high Monotonicity, matching expectations for random network assignment. Application of FERRET to both dimensionally reduced and genome-wide single-cell RNA sequencing datasets further demonstrated that the framework distinguishes methodological differences among GRN inference methods in ways that are consistent with their underlying statistical assumptions and biological design. Although these analyses were performed using three cancer datasets representing a limited number of cell types, the framework is general and may be applied to any collection of biologically related groups for which appropriate reference, in-group, and out-group networks can be defined.

FERRET complements existing benchmarking frameworks such as BEELINE by evaluating a different aspect of GRN inference performance. BEELINE measures the recovery of curated regulatory interactions using experimentally derived or simulated reference networks, providing an important assessment of edge-level accuracy. FERRET instead evaluates the reproducibility and biological consistency of inferred regulatory networks without relying on curated reference networks or simulated data. These two approaches therefore address distinct but complementary questions: whether a method recovers known regulatory interactions and whether it robustly captures biologically meaningful regulatory differences in real experimental data. Together, they provide a more comprehensive framework for benchmarking GRN inference methods than either approach alone.

Several limitations should be acknowledged. FERRET assumes that biologically meaningful in-group and out-group comparisons can be defined and that the selected groups differ in ways that are reflected in their underlying regulatory networks. The framework also depends on the choice of network similarity metric, although it is readily extensible to alternative measures of network similarity, including approaches based on spectral properties, modularity, or network geometry (41–43). Finally, biological interpretation remains dependent on current pathway annotations and the completeness of the literature used to evaluate enriched pathways.

The increasing diversity of GRN inference methods makes rigorous benchmarking both more important and more challenging. Because FERRET evaluates robustness and biological relevance directly from experimental data, it avoids many of the limitations associated with simulated datasets, curated regulatory networks, and the identification of true negative regulatory interactions. The framework is distributed as part of the NetZooR package together with benchmarking datasets from HTAN and CPTAC while remaining flexible enough to accommodate user-defined datasets, cell types, and similarity metrics. We anticipate that FERRET will provide a practical foundation for evaluating emerging GRN inference methods, including approaches based on multimodal data, spatial transcriptomics, temporal measurements, and increasingly sophisticated machine learning models.

## DATA AVAILABILITY

Data are available from Zenodo at https://zenodo.org/records/10854694 (CPTAC RCC), https://zenodo.org/records/10850845 (CPTAC GBM), and https://zenodo.org/records/14057537 (HTAN SCLC). FERRET is available from GitHub (https://github.com/QuackenbushLab/FERRET). Scripts for reproducing the analyses in this manuscript are also available from GitHub (https://github.com/QuackenbushLab/FERRET_paper_scripts).

## SUPPLEMENTARY DATA

Supplementary File 1: Supplementary Methods and References

Supplementary File 2: Supplementary Tables

Supplementary File 3: Supplementary Figures

## AUTHOR CONTRIBUTIONS

Tara Eicher: Conceptualization, Formal analysis, Methodology, Validation, Writing—original draft. John Quackenbush: Conceptualization, Writing—review & editing.

## ACKNOWLEDGEMENTS

We would like to thank Lauren Hsu, Chen Chen, Kimberly Glass, Dawn DeMeo, and Viola Fanfani for the suggestions regarding the development and validation of FERRET. We would further like to thank Anthony Gitter for his assistance in installing and running SINGE.

## FUNDING

This work was supported by the National Institutes of Health [R01HG011393, R35CA220523].

## CONFLICT OF INTEREST

None declared

## Notes

https://zenodo.org/records/10854694

https://zenodo.org/records/14057537

https://github.com/QuackenbushLab/FERRET

https://github.com/QuackenbushLab/FERRET_paper_scripts

### Competing Interest Statement

The authors have declared no competing interest.

