## Supplementary File 1 for "FERRET: Framework to Evaluate Robustness in Regulatory Networks Using Heterogeneous Cell Types"

### Supplementary Methods

#### Supplementary Equations

Supplementary Equation 1 is defined below, where *C* is the set of edge weight cutoffs defined by the user, $-\left( g \right)$ and $+\left( g \right)$ are defined as the subnetworks of $g$ with negatively weighted and positively weighted edges, respectively, $g_{i,c}$ is the subnetwork of $g$ with edges weighted above cutoff *c*, $d\left( g_{i},g_{j} \right)$ is defined as a similarity metric between GRNs *i* and *j*, and $sort\left( X,Y \right)$ refers to set *X* sorted in ascending order of the elements in set *Y*.

$${Trap}_{i}=\left( min\left( S_{in,i},S_{in,i-1} \right)-Min \right)\left( S_{out,i}-S_{out,i-1} \right)+\frac{\left( max\left( S_{in,i},S_{in,i-1} \right)-min\left( S_{in,i},S_{in,i-1} \right) \right)\left( S_{out,i}-S_{out,i-1} \right)}{2}$$

$$Ext=\left( S_{in,\left| C \right|}-Min \right)\left( Max-\max_{c\in C} \left( D_{out} \right) \right)$$

$$S_{in}=sort\left( D_{in},D_{out} \right),S_{out}=sort\left( D_{out},D_{out} \right)$$

$$Min=min\left( \min_{c\in C} \left( D_{in} \right),\min_{c\in C} \left( D_{out} \right) \right)$$

$$Max=max\left( \max_{c\in C} \left( D_{in} \right),\max_{c\in C} \left( D_{out} \right) \right)$$

$$D_{in}=\left\{ \bar{D}_{in,c} for all c\in C \right\},D_{out}=\left\{ \bar{D}_{out,c} for all c\in C \right\}$$

$$\bar{D}_{in,c}=\frac{1}{\left| G_{in} \right|}\sum_{g_{j}\in G_{in}} \left( d\left( -\left( g_{ref,c} \right),-\left( g_{j,c} \right) \right) \right)$$

$\bar{D}_{out,c}=\frac{1}{\left| G_{out} \right|}\sum_{g_{j}\in G_{out}} \left( d\left( -\left( g_{ref,c} \right),-\left( g_{j,c} \right) \right) \right)$	(1)

We note that the negative and positive edge subsetting are only applicable for GRN inference methods where negative edge weights represent inhibition and not low confidence, which can be specified by the user. Where negative edge weights represent low confidence, we instead re-scale the weights so that all are positive.

Supplementary Equation 2 describes the lower, upper, and average RAUC curves for consolidated results, redefining the *Min*, *Max*, $D_{in}$, and $D_{out}$ values used to calculate ${Trap}_{i}$ and $Ext$ for all reference-in-group-out-group configurations across all samples (defined by *Config*, where $N_{k}$ represents the number of similarity measures between reference and out-group GRNs within the range of bin *k*). We represent these as *Min’*, *Max’*, ${D_{in}^{'}}^{-}$and ${{D'}_{out}}^{-}$, $\bar{{D'}_{in}}$ and $\bar{{D'}_{out}}$, and ${D_{in}^{'}}^{+}$ and ${{D'}_{out}}^{+}$, respectively. We note that the vector of similarities for both the in-group and out-group will contain *b* values, where *b* is the number of bins (which we set to 10).

$$Min'=\min_{\left\{ ref, in,out \right\}\in Config,} \left( min\left( \min_{c\in C} \left( D_{in} \right),\min_{c\in C} \left( D_{out} \right) \right) \right)$$

$$Max'=\max_{\left\{ ref, in,out \right\}\in Config,} \left( max\left( \max_{c\in C} \left( D_{in} \right),\max_{c\in C} \left( D_{out} \right) \right) \right)$$

$Bins=\bigcup_{k=1}^{b} \left[ {Min}_{out}+\left( k-1 \right)\frac{{Max}_{out}-{Min}_{out}}{b} \right.,\left. {Min}_{out}+k\frac{{Max}_{out}-{Min}_{out}}{b} \right)$

$${D_{in}^{'}}^{-}=\bigcup_{k=1}^{b} \left\{ \min_{\zeta\in\left\{ 1...N_{k} \right\}} \left\{ D_{in} s.t. {Min}_{out}+\left( k-1 \right)\frac{{Max}_{out}-{Min}_{out}}{b}\leq D_{out}<{Min}_{out}+k\frac{{Max}_{out}-{Min}_{out}}{b} \right\} \right\}$$

$${{D'}_{out}}^{-}=\bigcup_{k=1}^{b} \left\{ \min_{\zeta\in\left\{ 1...N_{k} \right\}} \left\{ D_{out} s.t. {Min}_{out}+\left( k-1 \right)\frac{{Max}_{out}-{Min}_{out}}{b}\leq D_{out}<{Min}_{out}+k\frac{{Max}_{out}-{Min}_{out}}{b} \right\} \right\}$$

$$\bar{{D^{'}}_{in}}=\bigcup_{k=1}^{b} \left\{ \frac{1}{N_{k}}\sum_{\zeta=1}^{N_{k}} \left\{ D_{in} s.t. {Min}_{out}+\left( k-1 \right)\frac{{Max}_{out}-{Min}_{out}}{b}\leq D_{out}<{Min}_{out}+k\frac{{Max}_{out}-{Min}_{out}}{b} \right\} \right\}$$

$$\bar{{D'}_{out}}=\bigcup_{k=1}^{b} \left\{ \frac{1}{N_{k}}\sum_{\zeta=1}^{N_{k}} \left\{ D_{out} s.t. {Min}_{out}+\left( k-1 \right)\frac{{Max}_{out}-{Min}_{out}}{b}\leq D_{out}<{Min}_{out}+k\frac{{Max}_{out}-{Min}_{out}}{b} \right\} \right\}$$

$${D_{in}^{'}}^{+}=\bigcup_{k=1}^{b} \left\{ \max_{\zeta\in\left\{ 1...N_{k} \right\}} \left\{ D_{in} s.t. {Min}_{out}+\left( k-1 \right)\frac{{Max}_{out}-{Min}_{out}}{b}\leq D_{out}<{Min}_{out}+k\frac{{Max}_{out}-{Min}_{out}}{b} \right\} \right\}$$

$${{D'}_{out}}^{+}=\bigcup_{k=1}^{b} \left\{ \max_{\zeta\in\left\{ 1...N_{k} \right\}} \left\{ D_{out} s.t. {Min}_{out}+\left( k-1 \right)\frac{{Max}_{out}-{Min}_{out}}{b}\leq D_{out}<{Min}_{out}+k\frac{{Max}_{out}-{Min}_{out}}{b} \right\} \right\}$$

$${Max}_{out}=\max_{\left\{ ref, in,out \right\}\in Config,} \left( \max_{c\in C} \left( D_{out} \right) \right)$$

$${Min}_{out}=\min_{\left\{ ref, in,out \right\}\in Config,} \left( \min_{c\in C} \left( D_{out} \right) \right)$$

(2)

Supplementary Equations 3-5 define the similarity metrics, where $E\left( g \right)$ and $V\left( g \right)$ are the set of all edges and nodes in GRN *g*, respectively, and $w_{i}\left( a,b \right)$ is the score associated with edge $\left( a,b \right)$ in GRN $g_{i}$.

$Jaccard\left( g_{ref,c},g_{j,c} \right)=\frac{\sum_{\left( a,b \right)\in\left( E\left( g_{ref,c} \right)\cap E\left( g_{j,c} \right) \right)} \left( min\left( w_{ref,c}\left( a,b \right),w_{j,c}\left( a,b \right) \right) \right)}{\sum_{\left( a,b \right)\in\left( E\left( g_{ref,c} \right)\cup E\left( g_{j,c} \right) \right)} \left( max\left( w_{ref,c}\left( a,b \right),w_{j,c}\left( a,b \right) \right) \right)}$	(3)

$Indegree\left( g_{ref,c},g_{j,c} \right)=1-\sum_{b \in\left( V\left( g_{ref,c} \right)\cup V\left( g_{j,c} \right) \right)} \left| \frac{\sum_{a \in V\left( g_{ref,c} \right)} \left( w_{ref,c}\left( a,b \right) \right)}{2\sum_{\left( x,y \right) \in E\left( g_{ref,c} \right)} \left( w_{ref,c}\left( x,y \right) \right)}-\frac{\sum_{a \in V\left( g_{j,c} \right)} \left( w_{j,c}\left( a,b \right) \right)}{2\sum_{\left( x,y \right) \in E\left( g_{j,c} \right)} \left( w_{j,c}\left( x,y \right) \right)} \right|$	(4)

$Outdegree\left( g_{ref,c},g_{j,c} \right)=1-\sum_{a \in\left( V\left( g_{ref,c} \right)\cup V\left( g_{j,c} \right) \right)} \left| \frac{\sum_{b \in V\left( g_{ref,c} \right)} \left( w_{ref,c}\left( a,b \right) \right)}{2\sum_{\left( x,y \right) \in E\left( g_{ref,c} \right)} \left( w_{ref,c}\left( x,y \right) \right)}-\frac{\sum_{b \in V\left( g_{j,c} \right)} \left( w_{j,c}\left( a,b \right) \right)}{2\sum_{\left( x,y \right) \in E\left( g_{j,c} \right)} \left( w_{j,c}\left( x,y \right) \right)} \right|$	(5)

Supplementary Equations 6-7 describe the common in-group and out-group networks, where $\left( a,b \right)_{in}\in E\left( \cap\left( G_{in} \right) \right)$ and $\left( a,b \right)_{out}\in E\left( \cap\left( G_{out} \right) \right)$ and $sign\left( a,b \right)=1$ if the edge is activating and $sign\left( a,b \right)=-1$ if the edge is inhibitory. Networks where sign indicates confidence were rescaled to be positive only. Edges that are not present in a network were considered to have a weight of 0.

$w_{in}\left( a,b \right)=\left\{ \begin{aligned} \min_{i<\left| G_{in} \right|} \left( {sign}_{i}\left( a,b \right)*w_{i}\left( a,b \right) \right), \left| \sum_{i<\left| G_{in} \right|} \left( {sign}_{i}\left( a,b \right) \right) \right|=\left| G_{in} \right| \\ 0,\left| \sum_{i<\left| G_{in} \right|} \left( {sign}_{i}\left( a,b \right) \right) \right|<\left| G_{in} \right| \end{aligned} \right.$	(6)

$w_{out}\left( a,b \right)=\left\{ \begin{aligned} \min_{i<\left| G_{out} \right|} \left( {sign}_{i}\left( a,b \right)*w_{i}\left( a,b \right) \right), \left| \sum_{i<\left| G_{out} \right|} \left( {sign}_{i}\left( a,b \right) \right) \right|=\left| G_{out} \right| \\ 0,\left| \sum_{i<\left| G_{out} \right|} \left( {sign}_{i}\left( a,b \right) \right) \right|<\left| G_{out} \right| \end{aligned} \right.$	(7)

The differential network is defined in Supplementary Equation 8.

$w_{diff}\left( a,b \right)=\left\{ \begin{aligned} w_{in}\left( a,b \right)-w_{out}\left( a,b \right), {sign}_{in}\left( a,b \right)={sign}_{out}\left( a,b \right) \\ w_{in}\left( a,b \right), {sign}_{in}\left( a,b \right)\neq{sign}_{out}\left( a,b \right) \end{aligned} \right.$	(8)

The gene-gene differential network from dimensionality reduced data is defined in Equation 9 as ${g'}_{diff}$ with dimensionality $p \times p$ using the PCA loadings (*PC*, with dimensionality $p \times100$), as defined in Equation 10, where $g_{diff}$ is the adjacency matrix in the dimensionality-reduced space calculated as specified in Supplementary Equation 8, with dimensionality $100 \times100$.

${g'}_{diff}=PC\times g_{diff}\times{PC}^{T}$	(9)

Finally, we computed the differential targeting score *Tar* for each gene *b* as defined in Supplementary Equation 10. Recall that this can be computed using the differential network as defined in either Supplementary Equation 8 (for the whole genome space) or Supplementary Equation 9 (for the dimensionality reduced space).

${TAR}_{b}=\sum_{a s.t. \left( a,b \right)\in E({g^{'}}_{diff})} \left( w_{diff}\left( a,b \right) \right)$	(10)

#### Cell-type-Specific-Genes

We used the same cell-type-specific genes to define the pseudotime trajectory for CPTAC and HTAN data. These are defined in Supplementary Table 8, with the rationale for each cell type described below:

- **Macrophages:** We selected genes associated with naïve (*M0*), pro-inflammatory (*M1*), and anti-inflammatory (*M2*) subtypes (1) and defined a trajectory from M0 to M1 to M2.
- **Podocytes:** We selected genes associated with podocyte dedifferentiation (2,3) and defined a trajectory from differentiated to dedifferentiated.
- **Neurons and Interneurons:** We selected genes associated with increased proliferation of tumor cells (4) and defined a trajectory from lower to higher proliferation.
- **Oligodendrocytes:** We selected genes associated with myelin components that suggest stalled differentiation (5) and defined a trajectory from normal to stalled differentiation.
- **B cells:** We selected genes associated with B cell differentiation and defined a trajectory from precursor B cells to differentiated B cells.
- **T cells:** We selected genes associated with T cell differentiation and defined a trajectory from precursor T cells to differentiated T cells.
- **Endothelial cells:** We selected genes associated with endothelial cell differentiation and defined a trajectory from precursor endothelial cells to differentiated endothelial cells.
- **Fibroblasts cells:** We selected genes associated with fibrogenesis and defined a trajectory from lower to higher fibrogenesis.
- **NK cells:** We selected genes that mediate cytotoxicity and defined a trajectory from lower to higher mediation.

#### Gene Regulatory Network Inference Method Details

*SCENIC.* Single-Cell Regulatory Network Inference and Clustering (SCENIC), available as an R package, is a method for obtaining both comprehensive and cell-specific GRNs (10) from scRNA-seq. It finds candidate *regulons* (defined as groups of target genes co-regulated by a TF) using Gene Network Inference with Ensemble of Trees (GENIE3), a random forest based method for GRN inference (11). SCENIC then refines the set of regulons by finding regulatory modules that reflect known binding motifs derived from iRegulon (12). Finally, SCENIC determines the activity of each regulon across the cells. We evaluated SCENIC using the refined set of regulons and their associated correlation scores. Because SCENIC relies on a motif, it can only be run when the reference TF and gene labels are retained. For this reason, we only evaluated SCENIC using the whole-genome data and not the dimensionality reduced data.

*SCORPION.* Single-Cell Oriented Reconstruction of PANDA Individually Optimized gene regulatory Networks (SCORPION) (13), available as an R package, is a method for GRN inference from scRNA-seq data that extends the GRN inference method Passing Attributes between Networks for Data Assimilation (PANDA) (14) using a pseudo-bulking approach. PANDA is based on a message-passing framework that finds the consensus between TF motif binding sites, TF-specific protein-protein interactions (PPI), and target gene co-expression. We used TF motif binding sites and PPI derived from the Gene Regulatory Network Database (GRAND) (15). Like SCENIC, SCORPION depends on motifs defined using the reference gene labels. We therefore only evaluated SCORPION using the whole-genome data and not the dimensionality-reduced data. The scores assigned by SCORPION correspond to *z*-scaled confidence scores for each regulatory edge, where negative edge weights represent low confidence rather than inhibition.

*GRISLI.* Gene Regulatory Network Inference from scRNA-seq Data with Linear Differential Equations (GRISLI), available as a MATLAB script, is a GRN inference method for scRNA-seq data that is based on ordinary differential equations (ODE) computed using pseudotime (16). The magnitude of the score for each regulatory relationship corresponds to the impact of the source gene on the target gene, and the sign corresponds to the direction of regulation (inhibitory vs. activating).

*LEAP.* Lag-Based Expression Association for Pseudotime-Series (LEAP), available as an R package, is a GRN inference method for scRNA-seq data that is based on gene correlation over lag-based windows defined across pseudotime (17). The score for each regulatory edge represents the maximum absolute correlation between the genes across all time lags.

*PIDC.* Partial Information Decomposition and Context (PIDC), available as a Julia package, is a GRN inference method for scRNA-seq data that does not require pseudotime and is based on information gain on gene triplets (18). PIDC uses the concepts of *redundancy* (information about the expression of target gene *z* that can be obtained using either source genes *x* or *y*), *unique information* (information about the expression of *z* that can be obtained using only *x* or only *y*), and *synergistic information* (information about the expression of *z* that can be obtained using only the combination of both *x* and *y*) to infer regulatory relationships. The score for each regulatory edge can be positive only and represents a confidence value.

*scGeneRai.* Single-Cell Gene Regulatory Network Prediction by Explainable Artificial Intelligence (scGeneRai), available as a Python package, is a GRN inference method for scRNA-seq data that uses deep neural networks with layer-wise relevance propagation to infer a regulatory subnetwork for each individual target gene (19). The magnitude of the score for each regulatory relationship corresponds to the impact of the source gene on the target gene, and the sign corresponds to the direction of regulation (inhibitory vs. activating). Like SCENIC, scGeneRai infers cell-specific GRNs; it does so by first training a model using all cells and then applying it to each individual cell. We evaluated scGeneRai by averaging the scores for each edge across all GRNs for all cells.

*scSGL.* Single-Cell Graph Learning (scSGL), available as an R package, is a method that learns signed GRNs, where a negative score for an edge indicates that the source gene inhibits the target gene and a positive score indicates that the source gene activates the target gene (20). scSGL optimizes the GRN such that the total variation of gene expression is low with respect to the graph as reflected in the graph spectral domain. The magnitude of the score for each regulatory relationship corresponds to the impact of the source gene on the target gene.

*SINGE.* Single-Cell Inference of Networks using Granger Ensembles (SINGE), available as a MATLAB script and as a Docker image, is a GRN inference method for scRNA-seq data that relies on Granger causality inference across pseudotime (28). SINGE uses multiple window sizes with kernel smoothing to estimate the optimal lag. The magnitude of the score for each regulator relationship corresponds to the impact of the source gene on the target gene.

#### Biological Relevance of Pathway Categories

We determined biological relevance of pathways uncovered for each CPTAC and HTAN cell type by each GRN inference method by first manually assigning each pathway to a category denoting the central gene in the pathway (if specified in the pathway name) or the type of process (e.g., Amino Acid Metabolism). We then annotated each pathway category for each cell type. We marked broad categories (e.g., Infectious Disease, Cell Cycle, and Translation) or categories irrelevant to the cell type (e.g., Megakaryocytes, Germ Cells, or Cilia in T Cells) as “Not Searched”. We searched all remaining categories in PubMed using the name of the cell type and the name of the category, manually reviewing the top 10 results returned by PubMed when sorted by Best Match, with no additional filtering. PubMed searches were performed July 10, 2025. These were marked as either “No Support” or marked with the PubMed ID of the supporting paper. We included papers that described the activity of the pathway category in the cell type or the role of the pathway category in activation or cancer pathogenesis with respect to the cell type. We excluded papers that described a correlation between the pathway category and the cell type or that described the pathway category as being active in a subset of cells belonging to the cell type.

#### Definition of Bipartitification

Bipartitification is the graph transformation from an existing network $g_{i}$ to a new network $g_{i}'$ via three mappings.

We first define the following subsets of the original node and edge spaces in $g_{i}$ as follows:

$$V_{out}=\left\{ n\in V\left( g_{i} \right) :\exists m, \left( n,m \right)\in E\left( g_{i} \right) \right\}$$

$$V_{in}=\left\{ n\in V\left( g_{i} \right) :\exists m, \left( m,n \right)\in E\left( g_{i} \right) \right\}$$

Then we construct the node mappings, which are bijections from the original set of all nodes with out-degree and all nodes with in-degree to new node sets $U$ and $V$, respectively, as follows, where $n_{in}$ and $n_{out}$ are distinct formal copies of $n$:

$$B_{V}:V_{out}\to W, W=\left\{ n_{out} :n\in V_{out} \right\}, B_{W}\left( n \right)=n_{out}$$

$$B_{U}:V_{in}\to U, U=\left\{ n_{in} :n\in V_{in} \right\}, B_{U}\left( n \right)=n_{in}$$

Finally, we construct the edge mapping onto a new edge set $E$, which is a bijection, as follows:

$$B_{E}:E\left( g_{i} \right)\to E , E=\left\{ \left( a_{out},b_{in} \right) : \left( a,b \right)\in E\left( g_{i} \right) \right\},B_{E},B_{E}\left( \left( a,b \right) \right)=\left( a_{out},b_{in} \right)$$

We then define the new network $g_{i}'$ as follows:

$$V\left( g_{i}' \right)=U\cup W$$

$$E\left( g_{i}' \right)=E$$

We further define the weights of $E\left( g_{i}' \right)$ $w'$ as follows, where $w$ is the definition of weights in $E\left( g_{i} \right)$:

$$w'\left( a_{out},b_{in} \right)=w\left( a,b \right)$$

The following schematic illustrates bipartitification:

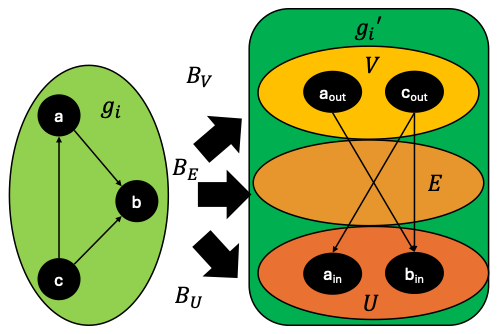

We have that *U* and *V* are disjoint copies of $V_{in}$ and $V_{out}$ by construction.

We also have that each edge in $E\left( g_{i}' \right)$ connects a node in *U* to a node in *V* by construction.

Because *U* and *V* are disjoint and each edge in $E\left( g_{i}' \right)$ connects a node in *U* to a node in *V*, $g_{i}'$ is bipartite.

#### Proof of Theorem 1 (Bipartitification Preserves the Jaccard Index in Networks with Positive Weights)

Let $g_{i}'$ and $g_{j}'$ be two bipartitified networks generated from directed networks $g_{i}$ and $g_{j}$, respectively, with at least one positive edge each, i.e., $\sum_{\left( a,b \right)\in\left( E\left( g_{i} \right)\cup E\left( g_{j} \right) \right)} \left( max\left( w_{i}\left( a,b \right),w_{j}\left( a,b \right) \right) \right)>0$

The Jaccard Index has been previously defined as follows, where $w_{i}\left( a,b \right)=0$ if $\left( a,b \right)\boldsymbol{\notin}E\left( g_{i} \right)$ and $w_{i}\left( a,b \right)>0$ if $\left( a,b \right)\boldsymbol{\in}E\left( g_{i} \right)$. We further define $w_{i}'\left( a_{out},b_{in} \right)=0$ if $\left( a,b \right)\boldsymbol{\notin}E\left( g_{i} \right)$.

$$Jaccard\left( g_{i},g_{j} \right)=\frac{\sum_{\left( a,b \right)\in\left( E\left( g_{i} \right)\cap E\left( g_{j} \right) \right)} \left( min\left( w_{i}\left( a,b \right),w_{j}\left( a,b \right) \right) \right)}{\sum_{\left( a,b \right)\in\left( E\left( g_{i} \right)\cup E\left( g_{j} \right) \right)} \left( max\left( w_{i}\left( a,b \right),w_{j}\left( a,b \right) \right) \right)}$$

Because $B_{E}\left( \left( a,b \right) \right)=\left( a_{out},b_{in} \right)$, we can write the following:

$$Jaccard\left( g_{i}',g_{j}' \right)=\frac{\sum_{\left( a,b \right)\in\left( E\left( g_{i} \right)\cap E\left( g_{j} \right) \right)} \left( min\left( w_{i}'\left( a_{out},b_{in} \right),w_{j}'\left( a_{out},b_{in} \right) \right) \right)}{\sum_{\left( a,b \right)\in\left( E\left( g_{i} \right)\cup E\left( g_{j} \right) \right)} \left( max\left( w_{i}'\left( a_{out},b_{in} \right),w_{j}'\left( a_{out},b_{in} \right) \right) \right)}$$

By definition, $w_{i}'\left( a_{out},b_{in} \right)=w_{i}\left( a,b \right)$ and $w_{j}'\left( a_{out},b_{in} \right)=w_{j}\left( a,b \right)$, yielding the following:

$$Jaccard\left( g_{i}',g_{j}' \right)=\frac{\sum_{\left( a,b \right)\in\left( E\left( g_{i} \right)\cap E\left( g_{j} \right) \right)} \left( min\left( w_{i}\left( a,b \right),w_{j}\left( a,b \right) \right) \right)}{\sum_{\left( a,b \right)\in\left( E\left( g_{i} \right)\cup E\left( g_{j} \right) \right)} \left( max\left( w_{i}\left( a,b \right),w_{j}\left( a,b \right) \right) \right)}$$

Therefore, $Jaccard\left( g_{i},g_{j} \right)=Jaccard\left( g_{i}',g_{j}' \right)$.

#### Proof of Theorem 2 (Bipartitification Preserves the In-Degree Index in Networks with Positive Weights)

Let $g_{i}'$ and $g_{j}'$ be two bipartitified networks generated from directed networks $g_{i}$ and $g_{j}$, respectively, with at least one positive edge each, i.e., $\sum_{\left( x,y \right) \in E\left( g_{i} \right)} \left( w_{i}\left( x,y \right) \right)>0$ and $\sum_{\left( x,y \right) \in E\left( g_{j} \right)} \left( w_{j}\left( x,y \right) \right)>0$.

The In-Degree Index has been previously defined as follows, where $w_{i}\left( a,b \right)=0$ if $\left( a,b \right)\boldsymbol{\notin}E\left( g_{i} \right)$ and $w_{j}\left( a,b \right)\boldsymbol{=}0$ if $\left( a,b \right)\boldsymbol{\notin}E\left( g_{j} \right)$. We further define $w_{i}'\left( a_{out},b_{in} \right)=0$ if $\left( a,b \right)\boldsymbol{\notin}E\left( g_{i} \right)$ or $\left( a_{out},b_{in} \right)\boldsymbol{\notin} E\left( g_{i}' \right)$ and $w_{j}'\left( a_{out},b_{in} \right)=0$ if $\left( a,b \right)\boldsymbol{\notin}E\left( g_{j} \right)$ or $\left( a_{out},b_{in} \right)\boldsymbol{\notin} E\left( g_{j}' \right)$.

$$Indegree\left( g_{i},g_{j} \right)=1-\sum_{b\in\left( V\left( g_{i} \right)\cup V\left( g_{j} \right) \right)} \left| \frac{\sum_{a \in V\left( g_{i} \right)} \left( w_{i}\left( a,b \right) \right)}{\sum_{\left( x,y \right) \in E\left( g_{i} \right)} \left( w_{i}\left( x,y \right) \right)}-\frac{\sum_{a \in V\left( g_{j} \right)} \left( w_{j}\left( a,b \right) \right)}{\sum_{\left( x,y \right) \in E\left( g_{j} \right)} \left( w_{j}\left( x,y \right) \right)} \right|$$

Because $B_{E}\left( \left( a,b \right) \right)=\left( a_{out},b_{in} \right)$, we can write the following:

$$Indegree\left( g_{i}',g_{j}' \right)=1-\sum_{l \in\left( V\left( g_{i}' \right)\cup V\left( g_{j}' \right) \right)} \left| \frac{\sum_{k \in V\left( g_{i}' \right)} \left( w_{i}'\left( k,l \right) \right)}{\sum_{\left( x,y \right) \in E\left( g_{i} \right)} \left( w_{i}'\left( x_{out},y_{in} \right) \right)}-\frac{\sum_{k \in V\left( g_{j}' \right)} \left( w_{j}'\left( k,l \right) \right)}{\sum_{\left( x,y \right) \in E\left( g_{j} \right)} \left( w_{j}'\left( x_{out},y_{in} \right) \right)} \right|$$

By definition, $w_{i}'\left( a_{out},b_{in} \right)=w_{i}\left( a,b \right)$ and $w_{j}'\left( a_{out},b_{in} \right)=w_{j}\left( a,b \right)$, yielding the following:

$$Indegree\left( g_{i}',g_{j}' \right)=1-\sum_{l \in\left( V\left( g_{i}' \right)\cup V\left( g_{j}' \right) \right)} \left| \frac{\sum_{k \in V\left( g_{i}' \right)} \left( w_{i}'\left( k,l \right) \right)}{\sum_{\left( x,y \right) \in E\left( g_{i} \right)} \left( w_{i}\left( x,y \right) \right)}-\frac{\sum_{k \in V\left( g_{j}' \right)} \left( w_{j}'\left( k,l \right) \right)}{\sum_{\left( x,y \right) \in E\left( g_{j} \right)} \left( w_{j}\left( x,y \right) \right)} \right|$$

Given that$B_{U}$ and $B_{V}$ are bijections, $\forall l\in\left( V\left( g_{i}' \right)\cup V\left( g_{j}' \right) \right)\exists b :b\in V\left( g_{i} \right)\cup V\left( g_{j} \right), l=b_{out}\vee l=b_{in}$ .

However, if $l=b_{out}$, then ${\forall k\in\left( V\left( g_{i}' \right)\cup V\left( g_{j}' \right) \right), w}_{i}'\left( k,l \right)=w_{j}'\left( k,l \right)=w_{i}'\left( k,b_{out} \right)=w_{j}'\left( k,b_{out} \right)=0$ because $b_{out}$ has no incoming edges following from the definition of bipartitification and therefore $\left( k,b_{out} \right)\boldsymbol{\notin} E\left( g_{i}' \right)\cup E\left( g_{j}' \right)$.

We may therefore simplify the In-Degree equation to the following:

$$Indegree\left( g_{i}',g_{j}' \right)=1-\sum_{b \in\left( V\left( g_{i} \right)\cup V\left( g_{j} \right) \right)} \left| \frac{\sum_{k \in V\left( g_{i}' \right)} \left( w_{i}'\left( k,b_{in} \right) \right)}{\sum_{\left( x,y \right) \in E\left( g_{i} \right)} \left( w_{i}\left( x,y \right) \right)}-\frac{\sum_{k\in V\left( g_{j}' \right)} \left( w_{j}'\left( k,b_{in} \right) \right)}{\sum_{\left( x,y \right) \in E\left( g_{j} \right)} \left( w_{j}\left( x,y \right) \right)} \right|$$

Similarly, we also have that $\forall k\in V\left( g_{i}' \right)\exists a :a\in V\left( g_{i} \right), k=a_{out}\vee k=a_{in}$ and $\forall k\in V\left( g_{j}' \right)\exists a :a\in V\left( g_{j} \right), k=a_{out}\vee k=a_{in}$.

However, if $k=a_{in}$, then ${\forall l\in\left( V\left( g_{i}' \right)\cup V\left( g_{j}' \right) \right), w}_{i}'\left( k,l \right)=w_{j}'\left( k,l \right)=w_{i}'\left( a_{in},l \right)=w_{j}'\left( a_{in},l \right)=0$ because $a_{in}$ has no outgoing edges following from the definition of bipartitification and therefore $\left( a_{in},l \right)\boldsymbol{\notin} E\left( g_{i}' \right)\cup E\left( g_{j}' \right)$.

We may therefore further simplify to the following:

$$Indegree\left( g_{i}',g_{j}' \right)=1-\sum_{b \in\left( V\left( g_{i} \right)\cup V\left( g_{j} \right) \right)} \left| \frac{\sum_{a \in V\left( g_{i} \right)} \left( w_{i}'\left( a_{out},b_{in} \right) \right)}{\sum_{\left( x,y \right) \in E\left( g_{i} \right)} \left( w_{i}\left( x,y \right) \right)}-\frac{\sum_{a\in V\left( g_{j} \right)} \left( w_{j}'\left( a_{out},b_{in} \right) \right)}{\sum_{\left( x,y \right) \in E\left( g_{j} \right)} \left( w_{j}\left( x,y \right) \right)} \right|$$

By definition, $w_{i}'\left( a_{out},b_{in} \right)=w_{i}\left( a,b \right)$ and $w_{j}'\left( a_{out},b_{in} \right)=w_{j}\left( a,b \right)$, yielding the following:

$$Indegree\left( g_{i}',g_{j}' \right)=1-\sum_{b \in\left( V\left( g_{i} \right)\cup V\left( g_{j} \right) \right)} \left| \frac{\sum_{a \in V\left( g_{i} \right)} \left( w_{i}\left( a,b \right) \right)}{\sum_{\left( x,y \right) \in E\left( g_{i} \right)} \left( w_{i}\left( x,y \right) \right)}-\frac{\sum_{a\in V\left( g_{j} \right)} \left( w_{j}\left( a,b \right) \right)}{\sum_{\left( x,y \right) \in E\left( g_{j} \right)} \left( w_{j}\left( x,y \right) \right)} \right|$$

Therefore, $Indegree\left( g_{i}',g_{j}' \right)=Indegree\left( g_{i},g_{j} \right)$.

#### Proof of Theorem 3 (Bipartitification Preserves the Out-Degree Index in Networks with Positive Weights)

Let $g_{i}'$ and $g_{j}'$ be two bipartitified networks generated from directed networks $g_{i}$ and $g_{j}$, respectively, with at least one positive edge each, i.e., $\sum_{\left( x,y \right) \in E\left( g_{i} \right)} \left( w_{i}\left( x,y \right) \right)>0$ and $\sum_{\left( x,y \right) \in E\left( g_{j} \right)} \left( w_{j}\left( x,y \right) \right)>0$.

The Out-Degree Index has been previously defined as follows, where $w_{i}\left( a,b \right)=0$ if $\left( a,b \right)\boldsymbol{\notin}E\left( g_{i} \right)$ and $w_{j}\left( a,b \right)=0$ if $\left( a,b \right)\boldsymbol{\notin}E\left( g_{j} \right)$. We further define $w_{i}'\left( a_{out},b_{in} \right)=0$ if $\left( a,b \right)\boldsymbol{\notin}E\left( g_{i} \right)$ or $\left( a_{out},b_{in} \right)\boldsymbol{\notin} E\left( g_{i}' \right)$ and $w_{j}'\left( a_{out},b_{in} \right)=0$ if $\left( a,b \right)\boldsymbol{\notin}E\left( g_{j} \right)$ or $\left( a_{out},b_{in} \right)\boldsymbol{\notin} E\left( g_{j}' \right)$.

$$Outdegree\left( g_{i},g_{j} \right)=1-\sum_{a\in\left( V\left( g_{i} \right)\cup V\left( g_{j} \right) \right)} \left| \frac{\sum_{b \in V\left( g_{i} \right)} \left( w_{i}\left( a,b \right) \right)}{\sum_{\left( x,y \right) \in E\left( g_{i} \right)} \left( w_{i}\left( x,y \right) \right)}-\frac{\sum_{b \in V\left( g_{j} \right)} \left( w_{j}\left( a,b \right) \right)}{\sum_{\left( x,y \right) \in E\left( g_{j} \right)} \left( w_{j}\left( x,y \right) \right)} \right|$$

Because $B_{E}\left( \left( a,b \right) \right)=\left( a_{out},b_{in} \right)$, we can write the following:

$$Outdegree\left( g_{i}',g_{j}' \right)=1-\sum_{k \in\left( V\left( g_{i}' \right)\cup V\left( g_{j}' \right) \right)} \left| \frac{\sum_{l \in V\left( g_{i}' \right)} \left( w_{i}'\left( k,l \right) \right)}{\sum_{\left( x,y \right) \in E\left( g_{i} \right)} \left( w_{i}'\left( x_{out},y_{in} \right) \right)}-\frac{\sum_{l \in V\left( g_{j}' \right)} \left( w_{j}'\left( k,l \right) \right)}{\sum_{\left( x,y \right) \in E\left( g_{j} \right)} \left( w_{j}'\left( x_{out},y_{in} \right) \right)} \right|$$

By definition, $w_{i}'\left( a_{out},b_{in} \right)=w_{i}\left( a,b \right)$ and $w_{j}'\left( a_{out},b_{in} \right)=w_{j}\left( a,b \right)$, yielding the following:

$$Outdegree\left( g_{i}',g_{j}' \right)=1-\sum_{k\in\left( V\left( g_{i}' \right)\cup V\left( g_{j}' \right) \right)} \left| \frac{\sum_{l \in V\left( g_{i}' \right)} \left( w_{i}'\left( k,l \right) \right)}{\sum_{\left( x,y \right) \in E\left( g_{i} \right)} \left( w_{i}\left( x,y \right) \right)}-\frac{\sum_{l \in V\left( g_{j}' \right)} \left( w_{j}'\left( k,l \right) \right)}{\sum_{\left( x,y \right) \in E\left( g_{j} \right)} \left( w_{j}\left( x,y \right) \right)} \right|$$

Given that$B_{U}$ and $B_{V}$ are bijections, $\forall k\in\left( V\left( g_{i}' \right)\cup V\left( g_{j}' \right) \right)\exists a :a\in V\left( g_{i} \right)\cup V\left( g_{j} \right), k=a_{out}\vee k=a_{in}$ .

However, if $k=a_{in}$, then ${\forall l\in\left( V\left( g_{i}' \right)\cup V\left( g_{j}' \right) \right), w}_{i}'\left( k,l \right)=w_{j}'\left( k,l \right)=w_{i}'\left( a_{in},l \right)=w_{j}'\left( a_{in},l \right)=0$ because $a_{in}$ has no outgoing edges following from the definition of bipartitification and therefore $\left( a_{in},l \right)\boldsymbol{\notin} E\left( g_{i}' \right)\cup E\left( g_{j}' \right)$.

We may therefore simplify the Out-Degree equation to the following:

$$Outdegree\left( g_{i}',g_{j}' \right)=1-\sum_{a\in\left( V\left( g_{i} \right)\cup V\left( g_{j} \right) \right)} \left| \frac{\sum_{l \in V\left( g_{i}' \right)} \left( w_{i}'\left( a_{out},l \right) \right)}{\sum_{\left( x,y \right) \in E\left( g_{i} \right)} \left( w_{i}\left( x,y \right) \right)}-\frac{\sum_{l \in V\left( g_{j}' \right)} \left( w_{j}'\left( a_{out},l \right) \right)}{\sum_{\left( x,y \right) \in E\left( g_{j} \right)} \left( w_{j}\left( x,y \right) \right)} \right|$$

Similarly, we also have that $\forall l\in V\left( g_{i}' \right)\exists b :b\in V\left( g_{i} \right), l=b_{out}\vee l=b_{in}$ and $\forall l\in V\left( g_{j}' \right)\exists b :b\in V\left( g_{j} \right), l=b_{out}\vee l=b_{in}$.

However, if $l=b_{out}$, then ${\forall k\in\left( V\left( g_{i}' \right)\cup V\left( g_{j}' \right) \right), w}_{i}'\left( k,l \right)=w_{j}'\left( k,l \right)=w_{i}'\left( k,b_{out} \right)=w_{j}'\left( k,b_{out} \right)=0$ because $b_{out}$ has no outgoing edges following from the definition of bipartitification and therefore $\left( k,b_{out} \right)\boldsymbol{\notin} E\left( g_{i}' \right)\cup E\left( g_{j}' \right)$.

We may therefore further simplify to the following:

$$Outdegree\left( g_{i}',g_{j}' \right)=1-\sum_{a \in\left( V\left( g_{i} \right)\cup V\left( g_{j} \right) \right)} \left| \frac{\sum_{b \in V\left( g_{i} \right)} \left( w_{i}'\left( a_{out},b_{in} \right) \right)}{\sum_{\left( x,y \right) \in E\left( g_{i} \right)} \left( w_{i}\left( x,y \right) \right)}-\frac{\sum_{b\in V\left( g_{j} \right)} \left( w_{j}'\left( a_{out},b_{in} \right) \right)}{\sum_{\left( x,y \right) \in E\left( g_{j} \right)} \left( w_{j}\left( x,y \right) \right)} \right|$$

By definition, $w_{i}'\left( a_{out},b_{in} \right)=w_{i}\left( a,b \right)$ and $w_{j}'\left( a_{out},b_{in} \right)=w_{j}\left( a,b \right)$, yielding the following:

$$Outdegree\left( g_{i}',g_{j}' \right)=1-\sum_{a \in\left( V\left( g_{i} \right)\cup V\left( g_{j} \right) \right)} \left| \frac{\sum_{b \in V\left( g_{i} \right)} \left( w_{i}\left( a,b \right) \right)}{\sum_{\left( x,y \right) \in E\left( g_{i} \right)} \left( w_{i}\left( x,y \right) \right)}-\frac{\sum_{b\in V\left( g_{j} \right)} \left( w_{j}\left( a,b \right) \right)}{\sum_{\left( x,y \right) \in E\left( g_{j} \right)} \left( w_{j}\left( x,y \right) \right)} \right|$$

Therefore, $Outdegree\left( g_{i}',g_{j}' \right)=Outdegree\left( g_{i},g_{j} \right)$.

### GRN Inference Methods Relying on Assumptions in Main Text Introduction

#### Random Forest Based Methods

- Huynh-Thu, V.A., Irrthum, A., Wehenkel, L. and Geurts, P. (2010) Inferring regulatory networks from expression data using tree-based methods. *PLoS One*, **5**.
- Shojaee, A. and Huang, S.C. (2023) Robust discovery of gene regulatory networks from single-cell gene expression data by Causal Inference Using Composition of Transactions. *Briefings in Bioinformatics*, **24**.

#### Gradient Boosting Based Methods

- Moerman, T., Aibar Santos, S., Bravo Gonzalez-Blas, C., Simm, J., Moreau, Y., Aerts, J. and Aerts, S. (2019) GRNBoost2 and Arboreto: efficient and scalable inference of gene regulatory networks. *Bioinformatics*, **35**, 2159-2161.
- Littman, R., Cheng, M., Wang, N., Peng, C. and Yang, X. (2023) SCING: Inference of robust, interpretable gene regulatory networks from single cell and spatial transcriptomics. *iScience*, **26**, 107124.

#### Neural Network Based Methods

- Chen, J., Cheong, C., Lan, L., Zhou, X., Liu, J., Lyu, A., Cheung, W.K. and Zhang, L. (2021) DeepDRIM: a deep neural network to reconstruct cell-type-specific gene regulatory network using single-cell RNA-seq data. *Briefings in Bioinformatics*, **22**.
- Keyl, P., Bischoff, P., Dernbach, G., Bockmayr, M., Fritz, R., Horst, D., Bluthgen, N., Montavon, G., Muller, K.R. and Klauschen, F. (2023) Single-cell gene regulatory network prediction by explainable AI. *Nucleic Acids Research*, **51**, e20.
- Luo, Q., Yu, Y. and Lan, X. (2022) SIGNET: single-cell RNA-seq-based gene regulatory network prediction using multiple-layer perceptron bagging. *Briefings in Bioinformatics*, **23**.
- Gan, Y., Hu, X., Zou, G., Yan, C. and Xu, G. (2022) Inferring Gene Regulatory Networks From Single-Cell Transcriptomic Data Using Bidirectional RNN. *Frontiers in Oncology*, **12**, 899825.
- Turki, T. and Taguchi, Y.-h. (2020) SCGRNs: Novel supervised inference of single-cell gene regulatory networks of complex diseases. *Computers in Biology and Medicine*, **118**.

#### Graph Laplacian Models

- Karaaslanli, A., Saha, S., Aviyente, S. and Maiti, T. (2022) scSGL: kernelized signed graph learning for single-cell gene regulatory network inference. *Bioinformatics*, **38**, 3011-3019.

#### Methods using Co-expression with Other Information

- Glass, K., Huttenhower, C., Quackenbush, J. and Yuan, G.-C. (2013) Passing Messages between Biological Networks to Refine Predicted Interactions. *PLoS ONE*, **8**.
- Weighill, D., M. Ben Guebila, K. Glass, J. Quackenbush and J. Platig (2022). Predicting genotype-specific gene regulatory networks. *Genome Research,* **32**(3), 524-533.
- Kuijjer, M. L., M. G. Tung, G. Yuan, J. Quackenbush and K. Glass (2019). Estimating Sample-Specific Regulatory Networks. *iScience*, **14**, 226-240.
- Osorio, D., Capasso, A., Eckhardt, S.G., Giri, U., Somma, A., Pitts, T.M., Lieu, C.H., Messersmith, W.A., Bagby, S.M., Singh, H. *et al.* (2024) Population-level comparisons of gene regulatory networks modeled on high-throughput single-cell transcriptomics data. *Nature Computational Science*, **4**, 237-250.
- Chen, C. and M. Padi (2024). Flexible modeling of regulatory networks improves transcription factor activity estimation. *npj Systems Biology and Applications,* **10**(1).

#### Causal Models from Temporal, Pseudotemporal, or RNA Velocity Data

- Aubin-Frankowski, P.C. and Vert, J.P. (2020) Gene regulation inference from single-cell RNA-seq data with linear differential equations and velocity inference. *Bioinformatics*, **36**, 4774-4780.
- Matsumoto, H., Kiryu, H., Furusawa, C., Ko, M.S.H., Ko, S.B.H., Gouda, N., Hayashi, T. and Nikaido, I. (2017) SCODE: an efficient regulatory network inference algorithm from single-cell RNA-Seq during differentiation. *Bioinformatics*, **33**, 2314-2321.
- Deshpande, A., Chu, L.F., Stewart, R. and Gitter, A. (2022) Network inference with Granger causality ensembles on single-cell transcriptomics. *Cell Reports*, **38**, 110333.
- Specht, A.T. and Li, J. (2017) LEAP: constructing gene co-expression networks for single-cell RNA-sequencing data using pseudotime ordering. *Bioinformatics*, **33**, 764-766.
- Papili Gao, N., Ud-Dean, S.M.M., Gandrillon, O. and Gunawan, R. (2018) SINCERITIES: inferring gene regulatory networks from time-stamped single cell transcriptional expression profiles. *Bioinformatics*, **34**, 258-266.
- Sanchez-Castillo, A., Vooijs, M. and Kampen, K.R. (2021) Linking Serine/Glycine Metabolism to Radiotherapy Resistance. *Cancers (Basel)*, **13**.
- Woodhouse, S., Piterman, N., Wintersteiger, C.M., Gottgens, B. and Fisher, J. (2018) SCNS: a graphical tool for reconstructing executable regulatory networks from single-cell genomic data. *BMC Systems Biology*, **12**, 59.
- Hossain, I., V. Fanfani, J. Fischer, J. Quackenbush and R. Burkholz (2023). Biologically Informed NeuralODEs for Genome-Wide Regulatory Dynamics. *BioRXiv.*

#### Information Gain Based Methods

- Chan, T.E., Stumpf, M.P.H. and Babtie, A.C. (2017) Gene Regulatory Network Inference from Single-Cell Data Using Multivariate Information Measures. *Cell Systems*, **5**, 251-267 e253.

#### Methods Using Transcription Factor Binding Motifs

- Glass, K., Huttenhower, C., Quackenbush, J. and Yuan, G.-C. (2013) Passing Messages between Biological Networks to Refine Predicted Interactions. *PLoS ONE*, **8**.
- Osorio, D., Capasso, A., Eckhardt, S.G., Giri, U., Somma, A., Pitts, T.M., Lieu, C.H., Messersmith, W.A., Bagby, S.M., Singh, H. *et al.* (2024) Population-level comparisons of gene regulatory networks modeled on high-throughput single-cell transcriptomics data. *Nature Computational Science*, **4**, 237-250.
- Aibar, S., Gonzalez-Blas, C.B., Moerman, T., Huynh-Thu, V.A., Imrichova, H., Hulselmans, G., Rambow, F., Marine, J.C., Geurts, P., Aerts, J. *et al.* (2017) SCENIC: single-cell regulatory network inference and clustering. *Nature Methods*, **14**, 1083-1086.
- Hossain, I., V. Fanfani, J. Fischer, J. Quackenbush and R. Burkholz (2023). Biologically Informed NeuralODEs for Genome-Wide Regulatory Dynamics. *BioRXiv.*
- Weighill, D., M. Ben Guebila, K. Glass, J. Quackenbush and J. Platig (2022). Predicting genotype-specific gene regulatory networks. *Genome Research,* **32**(3), 524-533.
- Chen, C. and M. Padi (2024). Flexible modeling of regulatory networks improves transcription factor activity estimation. *npj Systems Biology and Applications,* **10**(1).

#### Methods Using Protein-Protein Interactions

- Glass, K., Huttenhower, C., Quackenbush, J. and Yuan, G.-C. (2013) Passing Messages between Biological Networks to Refine Predicted Interactions. *PLoS ONE*, **8**.
- Osorio, D., Capasso, A., Eckhardt, S.G., Giri, U., Somma, A., Pitts, T.M., Lieu, C.H., Messersmith, W.A., Bagby, S.M., Singh, H. *et al.* (2024) Population-level comparisons of gene regulatory networks modeled on high-throughput single-cell transcriptomics data. *Nature Computational Science*, **4**, 237-250.

#### Methods Using Ground Truth Regulatory Networks

- Shojaee, A. and Huang, S.C. (2023) Robust discovery of gene regulatory networks from single-cell gene expression data by Causal Inference Using Composition of Transactions. *Briefings in Bioinformatics*, **24**.
- Chen, J., Cheong, C., Lan, L., Zhou, X., Liu, J., Lyu, A., Cheung, W.K. and Zhang, L. (2021) DeepDRIM: a deep neural network to reconstruct cell-type-specific gene regulatory network using single-cell RNA-seq data. *Briefings in Bioinformatics*, **22**.
- Gan, Y., Hu, X., Zou, G., Yan, C. and Xu, G. (2022) Inferring Gene Regulatory Networks From Single-Cell Transcriptomic Data Using Bidirectional RNN. *Frontiers in Oncology*, **12**, 899825.
- Wei, P.J., Guo, Z., Gao, Z., Ding, Z., Cao, R.F., Su, Y. and Zheng, C.H. (2024) Inference of gene regulatory networks based on directed graph convolutional networks. *Briefings in Bioinformatics*, **25**.
