## Supplementary Figure for "FERRET: Framework to Evaluate Robustness in Regulatory Networks Using Heterogeneous Cell Types"

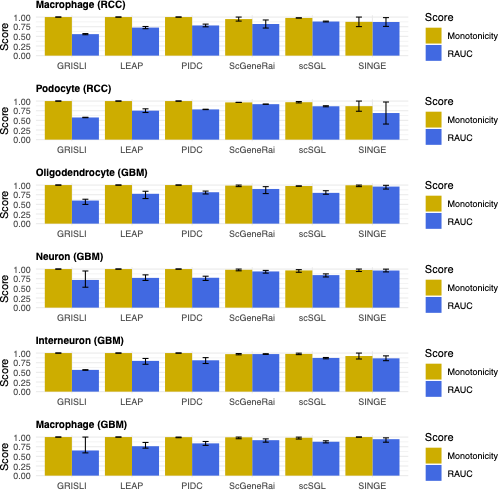

**Supplementary Fig. 1. RAUC and Monotonicity results for all methods evaluated on dimensionality reduced CPTAC data using In-Degree similarity.**

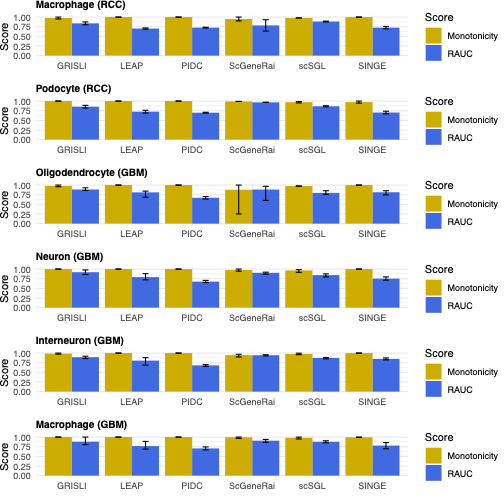

**Supplementary Fig. 2. RAUC and Monotonicity results for all methods evaluated on dimensionality reduced CPTAC data using Out-Degree similarity.**

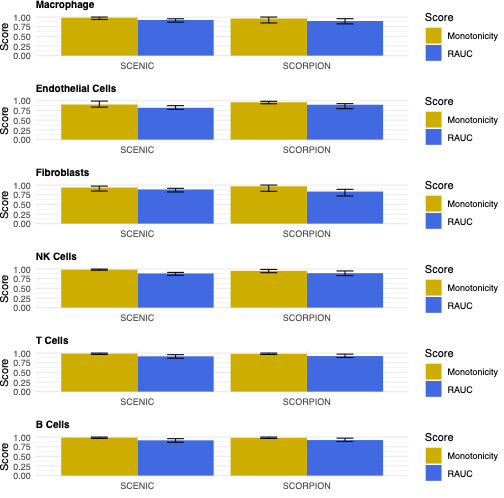

**Supplementary Fig. 3. RAUC and Monotonicity results for all methods evaluated on dimensionality reduced HTAN data using In-Degree similarity.**

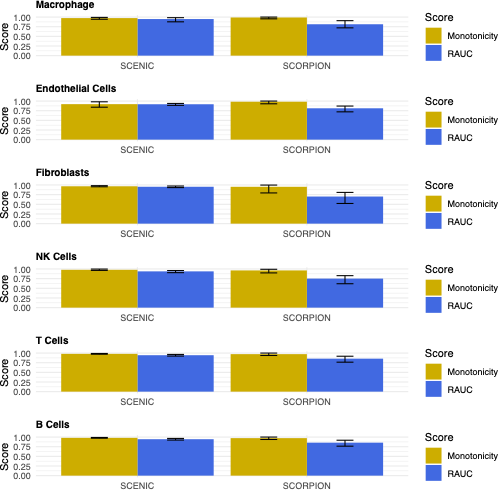

**Supplementary Fig. 4. RAUC and Monotonicity results for all methods evaluated on dimensionality reduced HTAN data using Out-Degree similarity.**
